# Interplay of ferroptotic and apoptotic cell death and its modulation by BH3-mimetics

**DOI:** 10.1101/2024.09.04.611207

**Authors:** Yun Qiu, Juliana A. Hüther, Bianca Wank, Antonia Rath, René Tykwe, Sabine Laschat, Marcus Conrad, Daniela Stöhr, Markus Rehm

## Abstract

Ferroptosis and apoptosis are widely considered to be independent cell death modalities. Ferroptotic cell death is a consequence of insufficient radical detoxification and progressive lipid peroxidation, which is counteracted by glutathione peroxidase 4 (GPX4). Apoptotic cell death can be triggered by a wide variety of stresses, including oxygen radicals, and can be suppressed by anti-apoptotic members of the BCL-2 protein family. Mitochondria are the main interaction site of BCL-2 family members and likewise a major source of oxygen radical stress. We therefore studied if ferroptosis and apoptosis might intersect and possibly interfere with one another. Indeed, cells dying from impaired GPX4 activity displayed hallmarks of both ferroptotic and apoptotic cell death, with the latter including (transient) membrane blebbing, submaximal cytochrome-c release and caspase activation. Targeting BCL-2, MCL-1 or BCL-XL with BH3-mimetics under conditions of moderate ferroptotic stress in many cases synergistically enhanced overall cell death and frequently skewed primarily ferroptotic into apoptotic outcomes. Surprisingly though, in other cases BH3-mimetics, most notably the BCL-XL inhibitor WEHI-539, counter-intuitively suppressed cell death and promoted cell survival following GPX4 inhibition. Further studies revealed that most BH3-mimetics possess previously undescribed antioxidant activities that counteract ferroptotic cell death at commonly employed concentration ranges. Our results therefore show that ferroptosis and apoptosis can intersect. We also show that combining ferroptotic stress with BH3-mimetics, context-dependently can either enhance and convert cell death outcomes between ferroptosis and apoptosis or can also suppress cell death by intrinsic antioxidant activities.

## Introduction

Ferroptosis and apoptosis are widely considered to be independent cell death modalities. Ferroptotic death results from excessive lipid peroxidation, which manifests when the cellular antioxidant systems are overwhelmed in their capacity to reduce (phospho)lipid hydroperoxides to their respective alcohols. In this process, the glutathione-dependent antioxidant system plays a key role with glutathione peroxidase 4 (GPX4) efficiently neutralizing phospholipid hydroperoxides at the expense of reduced glutathione (GSH). Cellular GSH levels are maintained by the activity of glutathione reductase or by *de novo* synthesis (Lu, 2013). For GSH synthesis, many cells rely on sufficient uptake of extracellular cystine, the oxidised and dimeric form of cysteine, through the cystine-glutamate antiporter system Xc-(Mandal et al., 2010). Indeed, cystine was identified as an essential component of common cell culture media already in the early days when developing tissue/cell culture systems (Eagle, 1955). Both GPX4 and system Xc- are pharmacological targets of synthetic ferroptosis-inducing agents, with RSL3 and erastin having been major hits in screens for compounds toxic to otherwise highly treatment resistant RAS-mutated cancer cells (Yang and Stockwell, 2008, Dolma et al., 2003). While RSL3 directly inhibits GPX4 by covalent interaction with the active site selenocysteine and was the lead for the development of related GPX4 inhibitors, erastin indirectly promotes ferroptosis by inhibiting system Xc- and cystine uptake, causing cellular GSH deprivation and GPX4 inactivation (Dixon et al., 2014, Yang et al., 2014). Ultimately, the membrane damage arising from widespread lipid peroxidation is thought to result in cell swelling and eventually cell rupture, whereby cellular content is released into the extracellular space. Whether specific molecular mechanisms are involved in the orchestration of this final stage of cell rupture during ferroptotic death is still being studied, with conflicting conclusions on the role of ninjurin-1 for plasma membrane permeabilisation (Hirata et al., 2023, Dondelinger et al., 2023).

In contrast to ferroptotic cell death, the primary endpoint of apoptosis is not cell swelling and lysis, but cell condensation and dismantling into membrane-enclosed apoptotic bodies or “blebs” (Kerr et al., 1972). Apoptosis is an active cellular response to external and internal stress conditions, which stands in contrast to ferroptosis as a consequence of perturbing steady-state cellular metabolism and redox fitness. In most scenarios of apoptotic cell death, mitochondrial outer membrane permeabilisation (MOMP) is a key decision point for triggering the execution of this form of cell death. MOMP is controlled by the BCL-2 protein family, with BAX and BAK constituting the major pore forming proteins and BCL-2, MCL-1 and BCL-XL acting as their antagonists (Czabotar and Garcia-Saez, 2023). A third group of BCL-2 family members are BH3-only proteins, of which the sensitizing BH3-only proteins act as neutralizers of BCL-2, MCL-1 and BCL-XL, thereby allowing BAX and/or BAK activation and oligomerisation to proceed. BH3-mimetics, small molecules that emulate the function of sensitizer BH3-only proteins, are now widely used in experimental settings to pharmacologically trigger MOMP (Czabotar and Garcia-Saez, 2023, Diepstraten et al., 2022). As a BCL-2 inhibitor, venetoclax has been approved for clinical use in a number of leukemias, while optimal translational avenues for inhibitors of MCL-1 and BCL-XL in single or combination treatments are currently being evaluated (Czabotar and Garcia-Saez, 2023, Diepstraten et al., 2022).

Mitochondria are a major site of membrane-associated redox signalling and likewise a source of cellular oxidative stress that could lead to ferroptotic cell death. At the same time, oxidative stress can sensitise to apoptosis (Redza-Dutordoir and Averill-Bates, 2016), where mitochondria are the central hub for the integration of apoptotic stress signals prior to triggering apoptosis execution. We therefore set out to study if both ferroptosis and apoptosis could possibly be coupled in response to GPX4 inhibition, and also if BH3-mimetics might modify cellular fates between survival and ferroptosis or apoptosis outcomes.

## Materials and methods

### Reagents and antibodies

S63845 was purchased from APExBIO Technology (Houston, TX, USA). ABT-199 was from Active Biochemicals Co. Limited (Hong Kong, China). A-1331852 was from TargetMol Chemicals Inc. (Boston, MA, USA), WEHI-539 hydrochloride was from MedChemExpress (Princeton, NJ, USA). Erastin was from Merck Chemicals GmbH (#329600, Darmstadt, Germany). DMSO and bovine serum albumin (BSA) were from Carl Roth (Karlsruhe, Germany). Q-VD-Oph (QVD), 4-hydroxytamoxifen and AZD5991 were from Selleckchem (Houston, TX, USA). Roswell Park Memorial Institute (RPMI) 1460 medium, Dulbecco’s Modified Eagle (DMEM) medium and sodium pyruvate were purchased from Thermo Fisher Scientific (Gibco, Waltham, MA, USA). Lipofectamine™ RNAiMAX transfection reagent, Hoechst 33342, C11 BODIPY 581/591, and MitoTracker CMXRos were purchased from Invitrogen (Carlsbad, CA, USA). Ferrostatin-1 (FER), (1S, 3R)-RSL3 (RSL3), glutathione reductase, fetal bovine serum, and propidium iodide (PI) were purchased from Sigma-Aldrich (Munich, Germany). Western blocking reagent and cOmplete protease inhibitor cocktail were from Roche Diagnostics (Mannheim, Germany). Paraformaldehyde was from Santa Cruz Biotechnology (Dallas, Texas, USA). 2,2-Diphenyl-1-picrylhydrazyl (DPPH) and STY-BODIPY were from Cayman Chemical Company (Ann Arbour, Michigan, USA). Fluoromount G was from Southern Biotechnology Associates Inc (Birmingham, Alabama, USA). 2,2’-Azobis(2-amidinopropane) dihydrochloride was from Sigma-Aldrich (St. Louis, Missouri, USA). Phosphatidylcholine chicken eggs was purchased from Lipoid (Ludwigshafen am Rhein, Rheinland-Pfalz, Germany). Diethylenetriaminepentaacetic acid, butylated hydroxytoluene, L-alpha-phosphatidylcholine from soybean, reduced glutathione, glutathione reductase from *Saccharomyces cerevisiae* were purchased from Sigma-Aldrich (St. Louis, Missouri, USA). SPLASH® LIPIDOMIX® was purchased from Avanti Polar Lipids (Alabaster, USA).

All antibodies used in immunoblotting were as follows: mouse monoclonal α-tubulin (1:5000, #3873), rabbit polyclonal β-Actin (1:1000, #4967), rabbit polyclonal caspase-3 (1:1000, #9662), mouse monoclonal BCL-2 (1:1000, #15071), rabbit monoclonal BCL-XL (1:1000, #2764), rabbit monoclonal MCL-1 (1:1000, #94296), rabbit polyclonal BAX (1:1000, #2772), rabbit polyclonal BAK (1:1000, #3814) and rabbit monoclonal Tom20 (1:1000, #42406) all from Cell Signaling Technology (Danvers, MA, USA). Rabbit monoclonal GPX4 (1:1000, ab125066) from Abcam (Cambridge, UK), mouse monoclonal PARP (1:1000, #556494, clone 4C10-5) and mouse monoclonal cytochrome-c (1:1000, #556433) from BD Pharmingen (Heidelberg, Germany). Goat anti-mouse IgG HRP-linked antibody (#115-035-062) and goat anti-rabbit IgG HRP-linked antibody (#111-035-144) were purchased from Dianova (Hamburg, Germany). The following antibodies were used in immunofluorescence staining: mouse monoclonal cytochrome-c (1:100) from Invitrogen (Carlsbad, CA, USA). Goat anti-rabbit IgG (H+L) Alexa Fluor 488 (#A-11008) and goat anti-mouse IgG (H+L) Alexa Fluor 647 (#A-21235) were from Invitrogen (Carlsbad, CA, USA).

### Cell culture

4-Hydroxytamoxifen-inducible *Gpx4^−/−^* immortalised mouse embryonic fibroblasts (Pfa1 cells) and HT1080M *Gpx4^−/−^* cells used in this study were described previously (Seiler et al., 2008, Doll et al., 2019). All other commercially available cell lines were used as authenticated materials (Microsynth, Göttingen, Germany). HT1080S and HT1080M are two strains of the HT1080 cell line that differ in their sensitivity towards RSL3. Both were authenticated as 100% HT1080 (**Supplemental Fig.1A,B**). All cell lines were grown in RPMI 1460 medium supplemented with 10% fetal bovine serum except HT-22 cells which were cultivated in DMEM supplemented with 10% fetal bovine serum and 1 mM sodium pyruvate. HT1080M *Gpx4^−/−^* cells were cultured in the presence of 2 µM FER. All cell lines were tested negative for mycoplasma contamination.

### Cell death assessment

Cells were seeded 24 h prior to stimulation. PI (1 µM) was used as a fluorescent indicator for cell death. Cells were pre-incubated with QVD for 30 min before the addition of the other reagents, where applicable. Cells were monitored in an IncuCyte S3 (Sartorius, Göttingen, Germany) and cell death was quantified using the IncuCyte analysis software, utilizing the cell-by-cell analysis module.

### Immunoblotting

Immunoblotting was performed as previously described (Stöhr et al., 2020). Cells were harvested by trypsinization and subsequently washed with cold phosphate-buffered saline (PBS, 2.67 mM KCl, 1.47 mM KH2PO4, 137.9 mM NaCl, 8.6 mM Na2HPO4, pH 7.4). Cells were centrifuged at 300 *g* for 5 min, with the subsequent removal of supernatants. Lysis buffer (150 mM NaCl, 1 mM EDTA, 20 mM TRIS, 1% (v/v) Triton X-100, pH 7.6) supplemented with 1× cOmplete protease inhibitor cocktail was added and the samples were incubated for 15 min on ice. Following centrifugation at 16,100 *g* for 15 min at 4°C, protein concentrations were determined by Bradford assay. Thereafter, 5 x Laemmli buffer (500 mM DTT, 10% (w/v) SDS, 25% (v/v) glycerol, 0.05% (w/v) bromophenol blue, 312.5 mM Tris-HCl) was added and the samples were incubated at 95°C for 5 min. Protein samples were separated using 4-12% Bis-Tris mini protein gels and transferred to a nitrocellulose membrane using an iBlot2 gel transfer device (Thermo Fisher Scientific, Waltham, MA, USA). Thereafter, the membrane was blocked by using a western blocking reagent for 1 h, and subsequently incubated with the primary and secondary antibodies.

### Cell fractionation

Cells were cultured in 6-well plates one day prior to stimulation. On the following day, after the designated stimulation period, the medium was aspirated. Then, 150 μl of permeabilisation buffer (210 mM D-Mannitol, 70 mM D-(+)-Sucrose, 10 mM HEPES, 5 mM Succinate, 0.5 mM EGTA, 100 µg/ml Digitonin, pH 7.2) were added to the centre of each well. The plate was gently rocked from side to side and from top to bottom for 45 seconds to ensure even distribution of the permeabilisation buffer. The supernatant from each well was collected and stored on ice (cytosolic fraction). Subsequently, 150 μl of lysis buffer containing 1x cOmplete protease inhibitor was added to the wells and pellet fractions were collected. Both fractions were then centrifuged at 13,000 *g* for 10 min at 4°C. The resulting supernatants were retained and protein concentrations were determined using the DC Protein Assay Kit II (Bio-Rad, Feldkirchen, Germany).

### Gene silencing with siRNA

Cells were seeded in 6-well plates and transfected 24 h later. For that, Lipofectamine® RNAiMAX Reagent and siRNA were diluted in Opti-MEM® medium according to the manufacturer’s instructions (Thermo Fisher Scientific, Waltham, MA, USA). The diluted siRNAs (10 nM) and Lipofectamine® RNAiMAX Reagent (10 nM) were mixed at a ratio of 1:1, and incubated at room temperature (RT) for 5 min. Thereafter, the solution was added drop by drop to each well. After 72 h, the transfection reagent was aspirated.

### C11 BODIPY 581/591 measurements

Cells were seeded one day prior to stimulation. The treatments were followed by incubation with C11 BODIPY 581/591 (1.5 µM) for 30 min at 37°C in the dark. Cells were harvested by trypsinisation, and centrifuged at 300 *g* for 5 min at RT. The cell pellet was resuspended in 100 µl PBS. For flow cytometric analysis, at least 10,000 events were acquired per sample using one of two flow cytometers: either a flow cytometer (CytoFLEX and CytExpert 2.4, Beckman Coulter), which utilizes a 488-nm laser paired with a 530/30 nm bandpass filter, or a flow cytometry (MACSQuant Analyzer 10, Miltenyi Biotec, Bergisch Gladbach, Germany), which is equipped with a 488-nm laser and a 525/50 nm bandpass filter for detection. Data from all experiments were analysed using the FlowJo software (Treestar).

### Lipidomic analysis

Pfa1 cells were seeded in 10 cm petri dishes (4 x10^5^ cells/dish). On the next day, cells were treated with RSL3, in the presence or absence of WEHI-539. Four hours later, cellular lipids were extracted using the methyl-tert-butyl ether (MTBE) method (Matyash et al., 2008). Briefly, cells were washed with antioxidant buffer (100 µM diethylenetriamine pentaacetate and 100 µM butylated hydroxytoluene in PBS, pH 7.4), and harvested by trypsinisation. Cells were centrifuged at 1500 *rpm* for 5 min at 4°C, cell pellets were washed with antioxidant buffer and centrifuged. The supernatant was removed and cell pellets were resuspended in 100 µl antioxidant buffer. Subsequently, samples were transferred into glass tubes containing a mixture of 1.5 ml ice-cold methanol, 3 ml chloroform, and 0.5 μl SPLASH^®^ LIPIDOMIX^®^ (Avanti Polar Lipids, Alabaster, USA). Samples were vortexed and incubated for 1 h at 4°C (Orbital shaker, 32 *rpm*). Phase separation was initiated by the addition of 1.25 ml water. The mixture was then vortexed and incubated for 10 min at 4°C. Subsequently, samples were centrifuged at 1500 *g* for 10 min at 4°C, resulting in the separation of the organic and aqueous phases. The organic phase was collected and the sample was re-extracted. The resulting lipid extracts were dried in a vacuum evaporator, dissolved in 120 µl isopropanol, and stored at -80°C before LC-MS analysis. Lipid extracts were supplemented with 0.01% (w/v) butylated hydroxytoluene and cooled on ice prior to the lipid extraction process.

Reversed phase liquid chromatography (RPLC) was performed on a Shimadzu ExionLC equipped with an AccucoreTM C18 column (150 x 2.1 mm; 2.6 µm, 80 Å, Thermo Fisher Scientific, Waltham, MA, USA). Oxidised lipids were separated by gradient elution with solvent A (acetonitrile/water, 1:1, v/v) and B (isopropanol/acetonitrile/water, 85:10:5, v/v), both containing 5 mM NH4HCO2 and 0.1% (v/v) formic acid. Separation was conducted at 50°C with a flow rate of 0.3 ml/min using the following gradient: 0-20 min, 10 to 86% B (curve 4); 20-22 min, 86 to 95% B; 22-26 min, 95% isocratic; 26-26.1 min, 95 to 10% B followed by 5 min re-equilibration at 10% B (Mishima et al., 2022). Mass spectrometry analysis was performed on a Sciex 7500 system equipped with an electrospray (ESI) source and operated in negative ion mode. Products were analysed in MRM mode, encompassing transitions from the precursor ion to the fragment ion, as described in **Supplemental Table 1**, with the following parameters: TEM 500°C, GS1 40, GS2 70, CUR 45, CAD 9, IS – 3,000 V. The area under the curve for the precursor mass to fragment mass was integrated and normalised by appropriate lipid species from SPLASH® LIPIDOMIX®. Normalised peak areas were further log-transformed (base 10) and auto-scaled in MetaboAnalyst online platform v5.0 (https://www.metaboanalyst.ca). Zero values were replaced by 0.2 times the minimum values detected for a given oxidised lipid within the samples. Oxidised lipids showing a significant difference (ANOVA, adjusted *P*-value (false discovery rate (FDR)), cut-off: 0.05) between samples were used for generating the heat maps. The heat maps were created in GraphPad Prism 8.0. The colour scheme corresponds to auto-scaled log fold change relative to the mean log value for the samples.

### Determination of GPX4 activity

Cells were seeded in 10 cm petri dishes and treated 24 h later with RSL3 in the presence or absence of WEHI-539 or FER for the indicated time points. The cells were washed with PBS, harvested, and then lysed in 100 µl of extraction buffer (1 mM EDTA, 150 mM KCl, 2 mM ß-mercaptoethanol, 0.1% (w/v) CHAPS, 100 mM KH2PO4/K2HPO4, pH 7.4). Cell lysates were subsequently centrifuged at 17,000 *g* and 4°C for 2 min. Then, 50 µl of protein supernatant was added to 1 ml reaction buffer (100 mM KH2PO4/K2HPO4, 5 mM EDTA, 5 mM glutathione, 0.1% peroxide-free Triton X-100, and 0.2 mM fresh NADPH). For the blank, 1 µl glutathione reductase was added in addition. Finally, 20 µl L-alpha-phosphatidylcholine hydroperoxide solution (25 µM) was added to initiate the reaction. GPX4 activity was measured by the absorbance decrease of NADPH at 340 nm.

For normalisation, the protein concentration of the samples was measured by Pierce™ 660 nm Protein Assay Kit using BSA as standard (Thermo Fisher Scientific, Waltham, MA, USA). The measurements were performed in a SpectraMax microplate reader (Molecular Device GmbH). The results were normalised to that of the DMSO control.

### FENIX assay

Egg phosphatidylcholine was weighed in a glass vial and dissolved in a minimal volume of chloroform. The solvent was evaporated under nitrogen, resulting in a thin film on the vial wall. The film was hydrated with PBS solution to produce a 20 mM lipid suspension. This lipid suspension underwent 10-15 freeze–thaw–sonication cycles, with each cycle involving freezing the vial in liquid nitrogen and thawing in a 37°C water bath. Finally, the lipid suspension was extruded 20–25 times using a mini extruder equipped with a 100 nm polycarbonate membrane (Avanti Polar Lipids, Alabaster, USA). Liposomes (from the above suspension) and STY-BODIPY (a 3 mM stock in DMSO) were diluted by PBS, resulting in the mastermix. The test compounds (2 µl) were added to a 96-well plate, followed by the addition of 295 µl mastermix. The plate was incubated for 10 min at 37°C in a plate reader. Subsequently, the plate was ejected from the plate reader, and 3 µl of 2,2’-Azobis (2-amidinopropane) dihydrochloride (AAPH, 1 mM) was added, followed by mixing for 5 min, resulting in final concentrations of 1 mM liposomes, 1 µM STY-BODIPY, 1 mM AAPH and the test compounds at the indicated concentrations. Fluorescence (λex = 488 nm, λem = 518 nm) was recorded every 10 min for the duration of the experiment. Data were transformed using the response factor of STY-BODIPY, followed by subtraction from the values of the DMSO control group in the absence of the radical initiator. Data were further analysed using GraphPad Prism 8.0.

### Immunofluorescence staining

Cells were seeded on sterilised glass coverslips one day prior to the experiment. MitoTracker CMXRos (200 nM) was added 30 min before the end of stimulation times. Afterwards, cells were washed two times with PBS and fixed with 4% paraformaldehyde (in PBS) for 10 min in the dark. Permeabilisation of the cells was carried out after three more washing steps with PBS using 0.1% (v/v) Triton X-100 (in PBS) for 10 min at RT and protected from light. Cells were washed once with PBS before unspecific binding sites were blocked with 4% BSA (in PBS) for 30 min at RT in the dark. After blocking, cells were incubated with the respective primary antibodies at RT for 1 h in the dark. Subsequently, cells were washed once with PBS and incubated with the respective fluorophore-labeled antibodies for 45 min at RT and protected from light. After incubation, cell nuclei were stained with Hoechst 33342 (1 µg/ml) for 10 min at RT in the dark. Subsequently, cells were washed three times with PBS before the coverslips were mounted on microscopy slides using Fluoromount G. Images of the probes were taken using a confocal microscope (Zeiss LSM710, Carl Zeiss, Jena, Germany), with an 40x/1.30 oil objective. Images were analysed using the CellProfiler software (Stirling et al., 2021). Masks were generated for cytoplasm and mitochondria and staining intensities were measured.

### Lactate dehydrogenase (LDH) assay

At specified time points, 100 μl of cell culture supernatant was collected. The remaining cells were lysed with 100 μl of 0.1% TritonX-100 solution in PBS. To determine the background signal, 100 μl of either cell culture media or 0.1% Triton X-100 solution in PBS was used. Subsequently, 100 μl of reaction master mix consisting of the components of the LDH reagent kit (Takara Bio Inc., Shiga, Otsu, Japan) (reagent A and reagent B in a ratio of 1:45) was added to each sample and incubated for a period of 15 to 30 min. The absorbance of the samples was measured at 492 nm by spectrophotometry. The results were normalised to the DMSO control.

### Live cell imaging

Cells were seeded one day prior to treatment in a four-compartment glass bottom dish. After stimulation with RSL3 in the presence or absence of QVD and/or FER (pre-incubation for 30 min), cells were imaged every 10 min for a total duration of 24 h using a Zeiss Cell Observer microscope (20x/0.80, Axio Observer.Z1/7, Carl Zeiss, Jena, Germany). Analysis of the cell morphology and cell fate was done manually in the Zen software.

### Cell death assessment using flow cytometry

Cells were seeded in 12-well plates one day prior to treatment. Cells were then stimulated with RSL3, in the presence of FER or WEHI-539. After the indicated times, cells were harvested by trypsinisation and centrifuged at 300 *g* for 5 min at RT. Cell pellets were resuspended in 100 µl PI solution (2 µg/ml in PBS) and incubated for 10 min at 37°C. The cell suspension was analysed using a flow cytometer (CytoFLEX and CytExpert 2.4, Beckman Coulter), and at least 10,000 events were analysed per sample. Data were analysed using FlowJo software (Treestar).

### Calculation of the coefficient of drug interaction (CDI)

CDIs were determined according to the Webb’s fractional product (Webb, 1963). CDI = E(AB) / (E(A) * E(B)), where E(A), E(B), and E(AB) represent the percentages of surviving cells after treatment with compound A, compound B, and the combination of A and B, respectively, relative to the control groups. CDI ≤ 0.9 indicates synergism, while CDI > 1.1 indicates antagonism.

### Long-term survival assay

Cells were subjected to continuous treatment with WEHI-539 either in the presence or absence of RSL3 over a 7-day period. Cells were split when cell densities reached 80-100%. Supernatants were replaced every two days. Cells were monitored and analysed using an IncuCyte S3 station, utilizing the cell-by-cell analysis module.

### Electrochemical methods

Differential pulse voltammetry (DPV) experiments were performed at RT and under Argon atmosphere. An AutolabPGSTAT204 potentiostat (Metrohm Autolab, Utrecht, The Netherlands) was used for all measurements. The experiment setup contained a three-electrode glass cell equipped with an AgCl-coated silver wire as pseudo reference electrode (RE), a Pt plate as counter electrode (CE) and a working electrode (WE) Au (vacuum deposited on glass slides over 3 nm of adhesion Cr layer; 30 nm Au layer) on S ∼ 0.5 cm^2^ slides. For all electrochemical measurements a 0.1 M solution of the supporting electrolyte Bu4NPF6 (Sigma Aldrich, electrochemical grade) in CH2Cl2 (Sigma Aldrich, HPLC grade, dry) was employed as received. All substrates were measured in a concentration of 0.3 mM in 0.1 M CH2Cl2 / Bu4NPF6 with a scan rate of 0.2 Vs-1 from 0 V to 2 V for the oxidation and from 0 V to -2 V for the reduction. The experiment solution was degassed by Argon before the measurements. The potentials were referenced to the formal potential of the Fc | Fc^+^ reference redox couple, measured in the same conditions as the analytes.

### Statistical analysis

GraphPad Prism 8.0 software (GraphPad Software, San Diego, CA, USA) was used for statistical analysis.

## Results

### GPX4 inhibition can trigger ferroptotic and apoptotic cell death

Ferroptosis and apoptosis are widely considered to be independent processes resulting in lytic or non-lytic cell death, respectively. Interestingly, we observed in two strains of HT1080 cells (**Supplemental Fig.1A,B**), the most widely used human cellular model system for ferroptosis studies, that GPX4 inhibition by RSL3 resulted in cell death that could be prevented partially by pan-caspase inhibitor QVD-OPH (**Fig.1A,B**). Suppression of cell death upon caspase inhibition was observed also when reducing RSL3 concentrations (**Supplemental Fig.1C-F**), or when inducing cell death by erastin, as an inhibitor of the cystine-glutamate antiporter system Xc^−^ (**Supplemental Fig.1G-J)**. Adding lipid ROS scavenger ferrostatin-1 fully prevented cell death, at least within the first 12 hours of treatment (**Fig.1A, B, Supplemental Fig.1C-F,H,J**), indicating that oxidative lipid damage appears to be the primary damage and prerequisite for a fraction of cells to develop a dependence on caspase activity to execute cell death. Data from surviving cells supported this further, since cells treated with RSL3 or erastin proliferated faster upon co-treatment with ferrostatin-1 when compared to QVD-OPH co-treatments (**Supplemental Fig.2A-D**). Still, caspase inhibition appears to clearly support survival and proliferation of GPX4-deficient cells, which rely on the continuous presence of ferrostatin-1, but which required substantially lower ferrostatin-1 concentrations in the presence of QVD-OPH (**Fig.1C, Supplemental Fig.2E**). Importantly, QVD-OPH did not prevent lipid peroxidation upon RSL3 treatment, as indicated by measuring C11 BODIPY conversion (**Fig.1D**), confirming that the cell-protective effect of caspase inhibition is downstream of lipid peroxidation arising from GPX4 inhibition.

**Figure 1:**
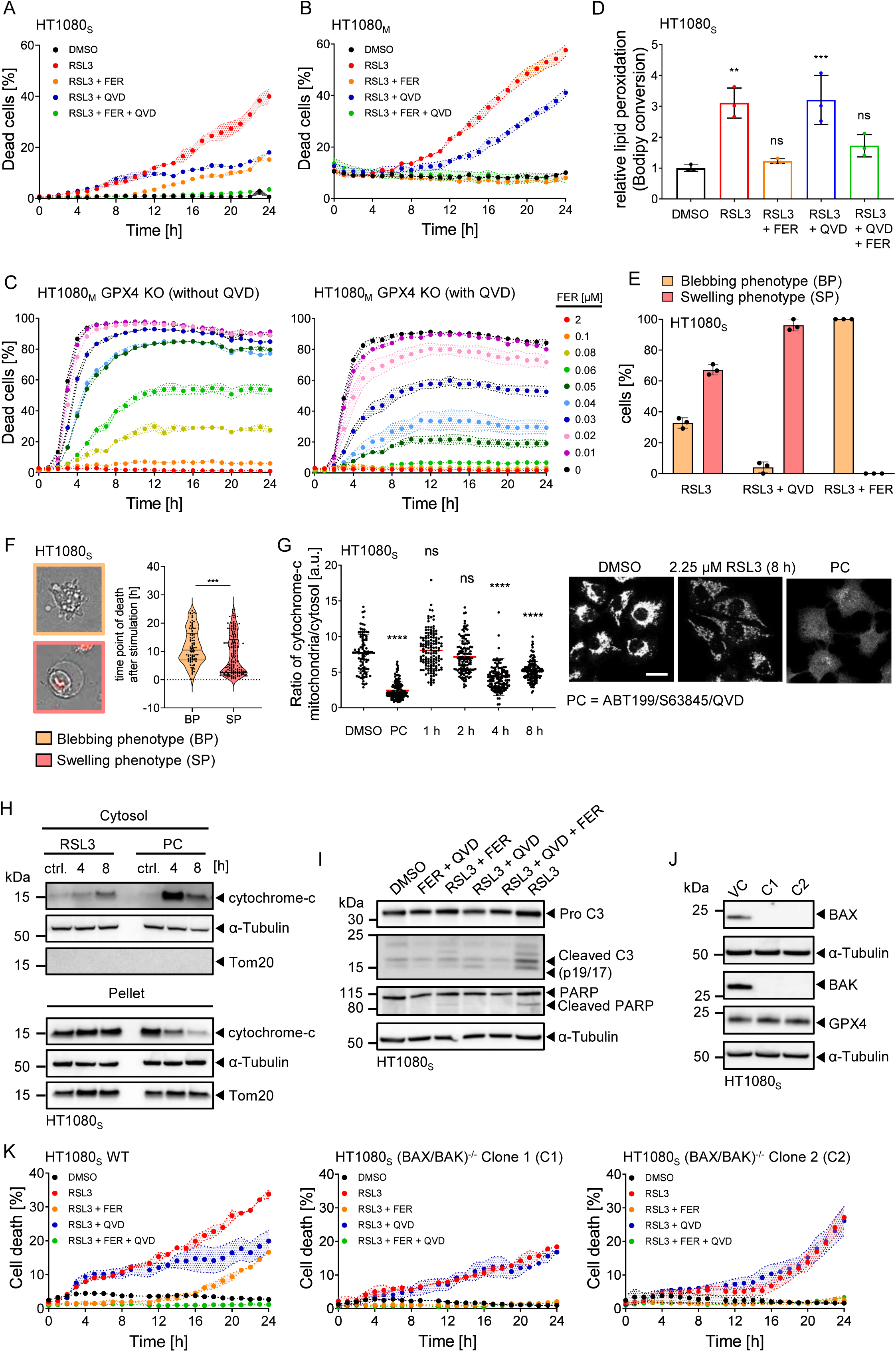
GPX4 inhibition can trigger ferroptotic and apoptotic cell death. **(A, B)** Quantification of cell death, calculated as percentage of PI-positive cells. Cells were stimulated with 4.5 µM RSL3 (HT1080S), 300 nM RSL3 (HT1080S), 50 µM QVD and 2 µM ferrostatin-1 (FER). Data are the mean ± range of technical duplicates **(A)** or mean ± SEM of technical triplicates **(B)**. Panels show one representative examples of at least three independent experiments. (C) Cells death was determined for cells cultured in presence or absence of 50 µM QVD and decreasing amounts of FER. Data are means ± SEM of technical triplicates of one representative example of three independent experiments. (D) Cells were stimulated with 4.5 µM RSL3 and analysed for lipid peroxidation (C11 BODIPY 581/591 conversion) via flow cytometry. Data show mean values ± SD of three independent experiments. Asterisks indicate statistical significance compared to the control group (* = p ≤ 0.05, ** = p ≤ 0.01; One-way Anova with Tukey’s post hoc test). (E) Cells were stimulated with 2.25 µM RSL3 and monitored at 10 min intervals via life-cell-imaging. Data are the mean ± SD of three independent experiments. (F) Data show median and quartiles of cells pooled from three independent experiments. Asterisks indicate statistical significance (*** = p ≤ 0.001; Mann-Whitney test). (G) Subcellular analysis of cytochrome-c redistribution. Cells were stimulated with 2.25 µM RSL3 for the indicated time points or with 10 µM ABT-199 + 10 µM S63845 + 50 µM QVD for 4 h as apoptotic positive control (PC). Fixed cells were immunostained for cytochrome-c and segmented based on MitoTracker signals. Scale bar in representative images, 20 µm. Data show means and scatter from one out of three representative experiments, with about 100 cells per condition. Asterisks indicate statistical significance compared to the control group (**** = p ≤ 0.0001; ns = not significant, One-way Anova with Tukey’s post hoc test). (H) Cells were stimulated with 4.5 µM RSL3 or 10 µM ABT-199 + 10 µM S63845 + 50 µM QVD (Positive ctrl. = PC) and fractioned into cytoplasm and pellet containing the mitochondria. One representative of three independent experiments is shown. (I) Immunoblotting of caspase-3 (C3) and PARP cleavage in HT1080S cells upon RSL3 (4.5 µM) treatment for 16 h, one representative of three independent experiments is shown. (J) Immunoblots of BAX, BAK and GPX4 expression in vector control cells (VC) and HT1080S (Bax/Bak)^-/-^ clones. (K) Quantification of cell death, calculated as percentage of PI-positive cells. Cells were stimulated with 2.25 µM RSL3, 50 µM QVD and 2 µM FER. Data show the mean ± range of technical duplicates of one representative example of four independent repeats.

Morphological studies of dying cells demonstrated that both cell swelling, a phenotype expected during the final stages of cells succumbing to ferroptosis, as well as cell shrinkage and membrane blebbing, indicative of apoptosis execution, could be observed in cells dying upon GPX4 inhibition (**Supplemental Movie 1**). A substantial portion of cells displayed, sometimes transient, membrane blebbing before inflating and taking up propidium iodide as a marker of impaired plasma membrane integrity. A quantification of these phenotypes in dying cells revealed that approximately one third of cells displayed membrane blebbing, transiently or terminally, following RSL3 treatment, and that this phenotype could be suppressed by caspase inhibition (**Fig.1E**). Cells with blebbing phenotypes tended to die later than cells that ruptured following cell swelling (**Fig.1F**). This indicates that within the intra-population heterogeneity of cell death phenotypes, features of apoptosis execution kinetically manifest somewhat later than cell death that is primarily ferroptotic. Taken together, these data support the notion that apoptosis signalling appears to contribute to cell death upon GPX4 inhibition, preferentially in fractions of cells that tend to die at later times.

Further evidence for engagement of apoptotic signalling upon GPX4 inhibition was obtained from analysing cytochrome-c distributions. RSL3 treatment resulted in cytochrome-c release into the cytosol, as evidenced in subcellular immunofluorescence analysis, yet less prominently than in positive controls of effective mitochondrial outer membrane permeabilization (**Fig.1G, Supplemental Figure 3**). Corresponding results were obtained by biochemical analysis of cytosolic cell extracts, in which cytochrome-c could be detected following GPX4 inhibition (**Fig.1H, Supplemental figure 4A-C**). This finding aligned with the observation that RSL3 treatment induced effector caspase-3 processing and, as proof of effector caspase activity, cleavage of PARP (**Fig.1I**). Apoptosis engagement was BAX/BAK dependent, since cells fully resistant to canonical induction of mitochondrial apoptosis (**Fig.1J, Supplemental Fig.4D**) were partially protected from RSL3, quantitatively identical to caspase inhibition by QVD-OPH co-treatment (**Fig.1K, Supplemental Fig.4E**).

Overall, these data therefore suggest that stress and cellular damage arising from GPX4 inhibition can induce cell death that proceeds with ferroptotic but also apoptotic features, wherein the latter requires the engagement of the mitochondrial apoptosis pathway.

### BH3-mimetics can synergistically enhance cell death induced by GPX4 inhibition

Mitochondrial apoptosis is governed by the BCL-2 protein family, whose anti-apoptotic family members can be targeted by highly specific pharmacological inhibitors (BH3-mimetics). BH3-mimetics are not only widely used in fundamental research but also are of substantial clinical interest as novel therapeutic candidates to combat otherwise non-responsive cancers in single or combination treatments (Diepstraten et al., 2022). Due to the indications for mitochondrial apoptotic engagement, we hypothesised that BH3-mimetics could promote cell death induced by GPX4 inhibition and possibly shift outcomes between ferroptotic and apoptotic cell death modalities.

We therefore treated HT1080S cells with concentration combinations of RSL3 and the clinically used BCL-2 inhibitor ABT-199, the BCL-XL inhibitor WEHI-539, or the MCL-1 inhibitor S63845. In all cases, combination treatments strongly and synergistically enhanced cell death (**Fig.2A-C**). Similar results were obtained for Pfa1 cells treated with RSL3 in combination with MCL-1 inhibitor S63845 or the alternative MCL-1 inhibitor AZD5991 (**Fig.2D**). In another ferroptosis-susceptible cell line, U87, AZD5991 in combination with RSL3 did not result in overt synergies at the measurement endpoint, but substantially accelerated cell death execution, likewise highlighting a positive interaction in this dual drug treatment (**Supplemental Fig.5**).

**Figure 2:**
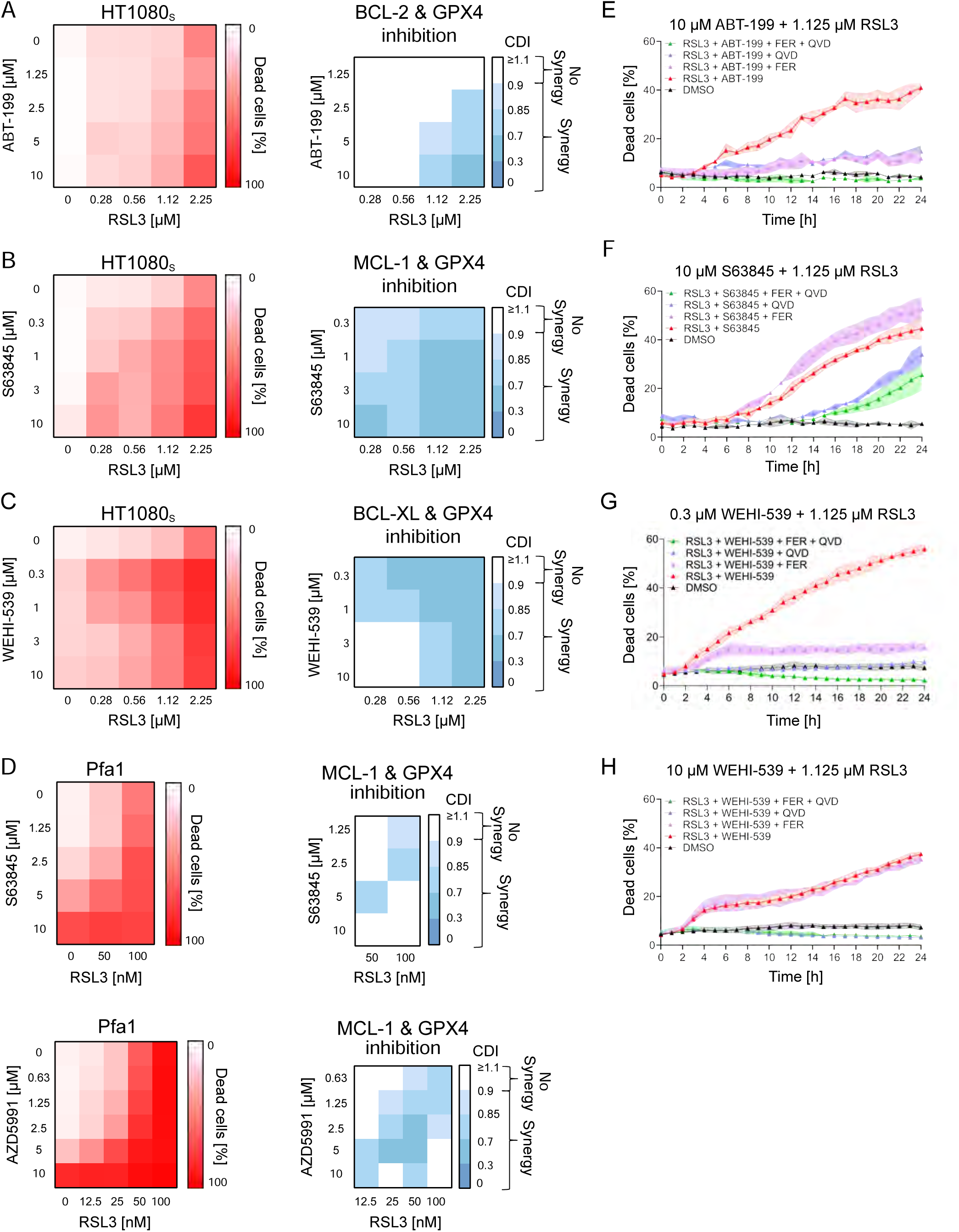
BH3-mimetics sensitise cells to RSL3-induced cell death. **(A-D)** Left heat map: Quantification of cell death, calculated as percentage of PI-positive cells. Cells were stimulated with the indicated concentrations of RSL3, the BCL-2 inhibitor ABT-199, the MCL-1 inhibitors S63845 or AZD5991, or the BCL-XL inhibitor WEHI-539 for 24 h. Data shown are means of two or three technical replicates. Right heat map: The coefficient of drug interaction (CDI) was calculated as Webb’s fractional product. **(E-H)** Cell death was determined by PI uptake. Cells were treated as indicated, in the presence or absence of ferrostatin-1 (FER, 2 µM) or QVD (50 µM). Data are mean ± SD of three technical replicates. Panels show one representative example of at least 2 independent experiments.

We next investigated the influence of BH3-mimetics on the type of cell death induced by RSL3. Synergistic cell death of HT1080S cells exposed to RSL3 and ABT-199 could be partially prevented by both QVD-OPH and ferrostatin-1 (**Fig.2A,E**), demonstrating both ferroptotic and apoptotic contributions to cell death. In contrast, the synergistic cell death upon combining RSL3 with the MCL-1 inhibitor S63845 predominantly induced cell death that was no longer prevented by ferrostatin-1, but partially by QVD-OPH, indicating a shift towards apoptosis as the primary terminal cell fate (**Fig.2F**). The synergistic induction of cell death by the combination of RSL3 and WEHI-539 was most pronounced at lower amounts of the BCL-XL inhibitor. At these concentrations, cells were partially protected by ferrostatin-1 but fully rescued by QVD-OPH (**Fig.2G**). At higher concentrations of WEHI-539, ferrostatin-1 lost its protective function and cell death apparently fully depended on caspase activity (**Fig.2H**).

Together these data reveal that the addition of BH3-mimetics can synergistically enhance RSL3-induced cell death and furthermore that this is accompanied by shifting the cell death modality towards apoptosis.

### BH3-mimetics can suppress cell death induced by GPX4 inhibition

During our analyses on the cell death enhancing effects of BH3-mimetics, we noted that the BCL-XL inhibitor WEHI-539 appeared to reduce peak cell death in HT1080S cells when applied in concentrations of 1 µM and higher (see **Fig.2C**). We therefore expanded our studies with this BH3-mimetic to additional cell lines and conditions. To our surprise, WEHI-539 very potently suppressed RSL3-induced death in Pfa1 and U87 cells, as evidenced by reduced PI uptake (**Fig.3A,B**) and by reduced LDH release (**Supplemental Fig.6**). A screen across a wider range of concentration combinations confirmed WEHI-539 to dose-dependently and potently antagonize RSL3-induced cell death in these cell lines (**Fig.3C**). Comparable results were obtained when replacing RSL3 with alternative GPX4 inhibitors, ML-162 and ML-210 (**Fig.3D**). A suppression of cell death upon GPX4 inhibition was observed likewise when using A-1331852 as an alternative BCL-XL inhibitor in Pfa1 and U87 cells (**Fig.3E**) and when combining the MCL-1 inhibitor S63845 with RSL3 in U87 cells (**Fig.3F**). WEHI-539 also suppressed cell death in HT-22 cells (**Fig.3G**), a hippocampal neuronal cell line frequently used to study ferroptosis susceptibilities (Chu et al., 2020, Neitemeier et al., 2017), as did ABT-199 in U87 cells (**Fig.3H**).

**Figure 3:**
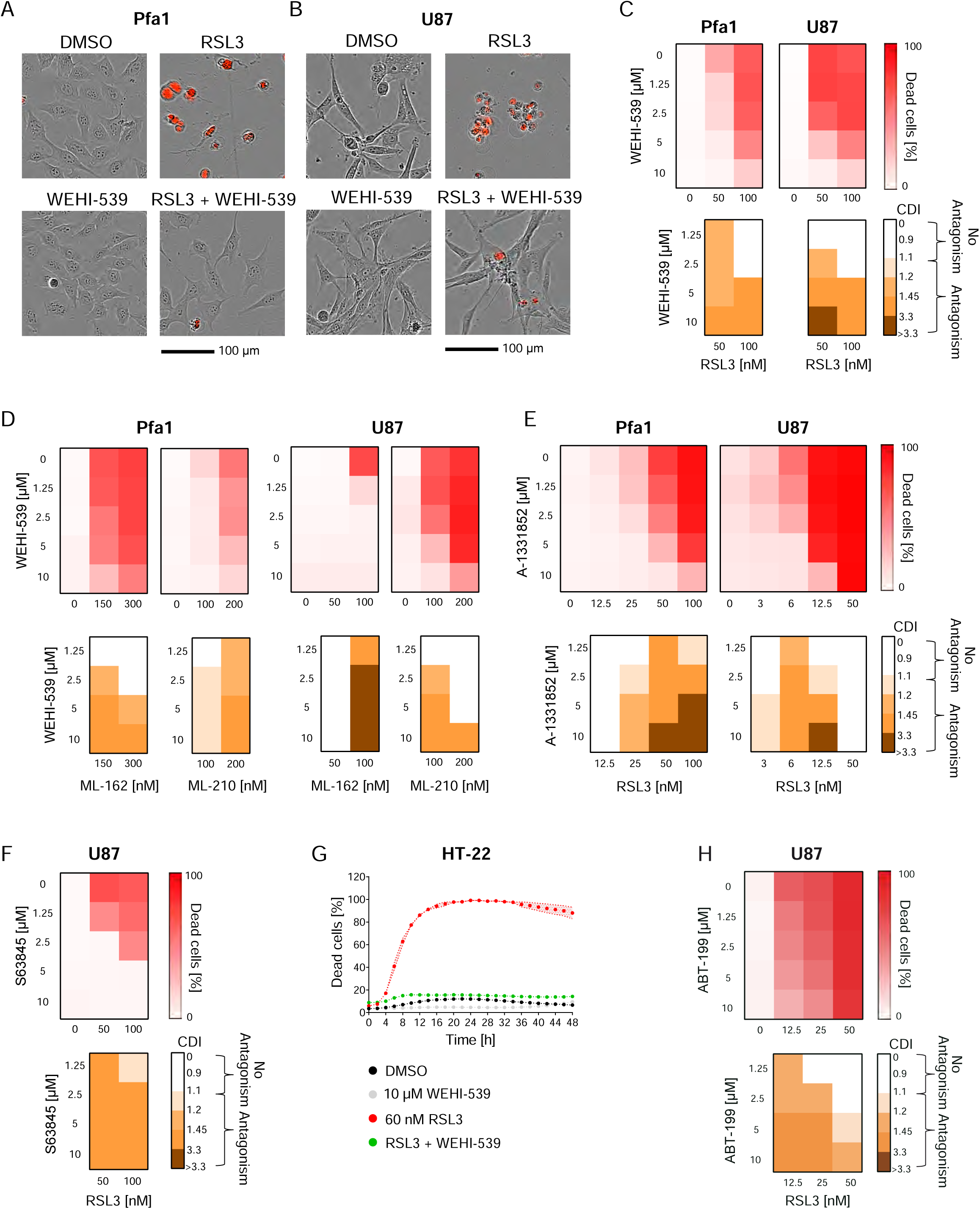
BH3-mimetics can suppress cell death induced by GPX4 inhibition. **(A, B)** Brightfield images with PI fluorescence overlays show cells treated for 24 h with either WEHI-539 (10 µM), RSL3 (100 nM), or both combined. **(C-F, H)** Cells were treated for 24 h with the indicated concentrations of GPX4 inhibitors (RSL3, ML-162, ML-210) in combination with BCL-XL inhibitors (WEHI-539, A-1331852), or the MCL-1 inhibitor S63845, or the BCL-2 inhibitor ABT-199. Cell death was determined by PI uptake. Antagonism was determined from the coefficient of drug interaction CDI by the Webb’s fractional product. Data represent the mean of two or three technical replicates, panels show one representative of at least two independent experiments **(G)** Quantification of cell death, calculated as percentage of PI-positive cells. Data are the mean ± SEM of technical triplicates. One representative experiment of four independent experiments is shown.

These results demonstrate a striking contrast to the previous conditions, where synergistic cell death induction had been observed for combination treatments with RSL3 and BH3-mimetics, demonstrating that BH3-mimetics can context-dependently suppress cell death in response to GPX4 inhibition.

### Cell death commitment upon GPX4 inhibition and dependencies on WEHI-539 for cell survival

Based on WEHI-539 as a representative BH3-mimetic, we next further studied the potency of ferroptosis antagonism upon GPX4 inhibition as well as effects on cell death susceptibility upon long-term exposure. Both Pfa1 and U87 cells regained proliferative capacity in continuous presence of RSL3/WEHI-539 after approximately 24 h of treatment, with proliferation rates similar to those of controls (**Fig.4A**). WEHI-539 therefore not only suppresses cell death but also permits cells to overcome or adapt to RSL3-induced stress, so that proliferation resumes. To understand if differences exist between cell protection by WEHI-539 and ferrostatin-1 in combination treatments with RSL3, drug-containing medium was aspirated and replaced with fresh growth medium after 24 h of incubation (**Fig.4B**). U87 control cells resisted 24 h RSL3 treatment in presence of either ferrostatin-1 or WEHI-539 during the combination treatment (**Fig.4C**), corresponding to the screening analyses in **Fig.3**. However, U87 cells died upon removal of WEHI-539 but not upon removal of ferrostatin-1 from the medium when RSL3 was washed out (**Fig.4D**). In Pfa1 cells, only a comparably small fraction of cells died upon washout, yet also here the removal of WEHI-539 was more likely to result in cell death than removing ferrostatin-1 (**Supplemental Fig.7A,B**). To also clarify if cells can adapt to RSL3-induced stress in presence of WEHI-539 in the long term, we continuously treated U87 and Pfa1 cells for 2-3 weeks with either WEHI-539 or RSL3/WEHI-539, followed by full drug washout or selective WEHI-539 or RSL3 washout. Both cell lines retained RSL3 responsiveness regardless of whether they were cultured in WEHI-539- or RSL3/WEHI-539-containing medium, yet RSL3 responsiveness was diminished in Pfa1 cells upon full RSL3/WEHI-539 drug washout, as only a fraction of cells committed cell death in this setting (**Fig.4E**). Overall, these results indicate that cells grown in medium containing RSL3/WEHI-539 for prolonged periods develop a dependency on the continued presence of WEHI-539 extending to times after which RSL3 has already been removed.

**Figure 4:**
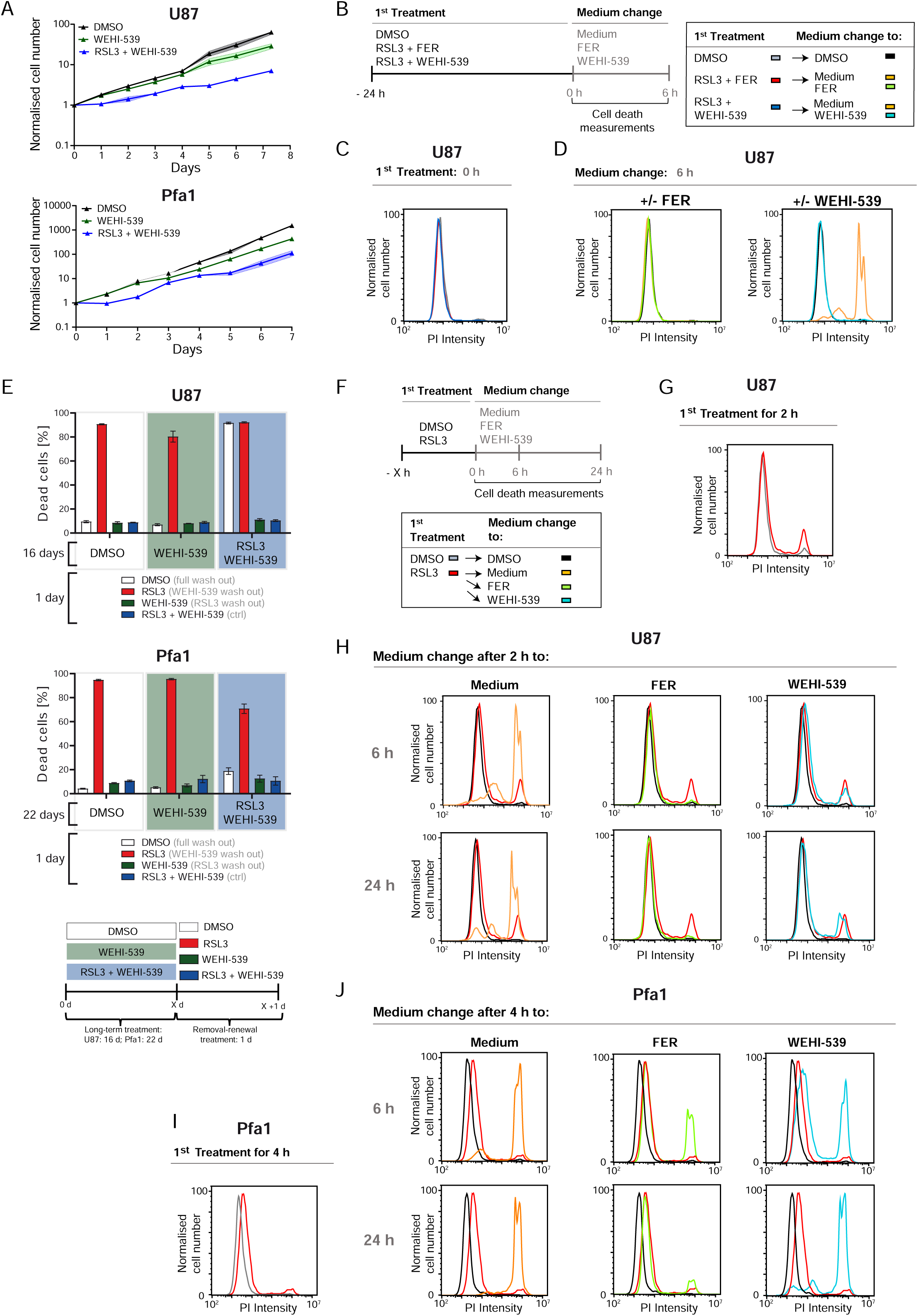
Cell death commitment upon GPX4 inhibition and dependencies on WEHI-539 for cell survival. **(A)** Cells were treated with WEHI-539 (10 µM) and proliferation was observed for 7 days, both in the presence and absence of RSL3 (100 nM). Supernatants were changed every second day. Data are mean ± SD of technical triplicates from one representative of 2 independent experiments. **(B-D)** The scheme **(B)** illustrates the 24 h treatment schedule with RSL3 (100 nM) and WEHI-539 (10 µM) or ferrostatin-1 (FER, 2 µM), followed by PBS washing. Cell death was measured by PI uptake via flow cytometry after 6 h. Data are from one representative of three independent experiments. **(E)** Cells were subjected to prolonged treatment with RSL3 (100 nM) and WEHI-539 (10 µM) for 16 days (U87 cells) or 22 days (Pfa1 cells), then washed with PBS and re-exposed as indicated. Cell death was determined by PI uptake. Data shown are mean ± SD of technical triplicates from one representative of two (Pfa1) or three (U87) independent experiments. **(F-J)** Scheme **(F)** displays the treatment schedule. Cells received a 1st treatment with DMSO or RSL3 (100 nM) for 2 h **(G)** or 4 h **(I)**. Cells were then washed and exposed to either FER or WEHI-539 **(H, J)**. Cell death was measured by PI uptake after the first treatment as well as 6 h and 24 h after the medium change. One representative of two (F-J) or three (B-D) independent experiments is shown.

To understand whether a “point of no return” in cell death commitment upon RSL3 exposure can be identified, we next treated U87 cells with RSL3, followed by time-shifted addition of ferrostatin-1 or WEHI-539 (**Fig.4F**). A pre-treatment of 2 h was sufficient for the first U87 cells to die from exposure to RSL3 (**Fig.4G**). Replacing the supernatant with regular growth medium failed to rescue the majority of cells from dying, whereas both ferrostatin-1 and WEHI-539 prevented cell death execution for at least 24 h of follow up observation (**Fig.4H**). In Pfa1 cells, cell death upon addition of ferrostatin-1 or WEHI-539 after a 4 h RSL3 pre-treatment was effectively suppressed by ferrostatin-1. Compared to the control, also WEHI-539 protected Pfa1 cells, yet less efficiently than ferrostatin-1 (**Fig.4I,J**). Reducing the pre-treatment duration to 2 h in Pfa1 cells substantially enhanced the capability of WEHI-539 to avert cell death (**Supplemental Fig.8A,B**). These data thus show that the BH3-mimetic WEHI-539 can exert its protective effect downstream of the initial GPX4 inhibition and that a point of no return has not yet been (entirely) passed 2-4 h later, since WEHI-539 as well as ferrostatin-1 can still rescue these cells.

### BH3 mimetics exhibit off-target activities that can suppress ferroptotic cell death

Next, we evaluated if off-target effects could be responsible for the anti-ferroptotic activity of BH3-mimetics. First, we studied if direct neutralization of RSL3 by WEHI-539 can be ruled out. In both U87 and Pfa-1 cells, WEHI-539 failed to prevent RSL3 from inhibiting GPX4 activity, as did ferrostatin-1 (**Fig.5A**). Instead of pharmacologically inhibiting GPX4, we also assessed if cells depleted of GPX4 by conditional tamoxifen-induced *Gpx4* targeting (Seiler et al., 2008) could be rescued by WEHI-539. Indeed, WEHI-539 effectively prevented cell death upon GPX4 loss (**Fig.5B**). Strikingly, depleting cells of BCL-XL expression did not prevent WEHI-539 from suppressing RSL3-induced cell death (**Fig.5C**). Similarly, the MCL-1 inhibitor S63845 could still prevent RSL3-induced cell death when the expression of its primary pharmacological target was suppressed (**Fig.5D**). Overall, this indicates that BH3-mimetics prevent ferroptotic cell death through off-target activities downstream of GPX4 inhibition.

**Figure 5:**
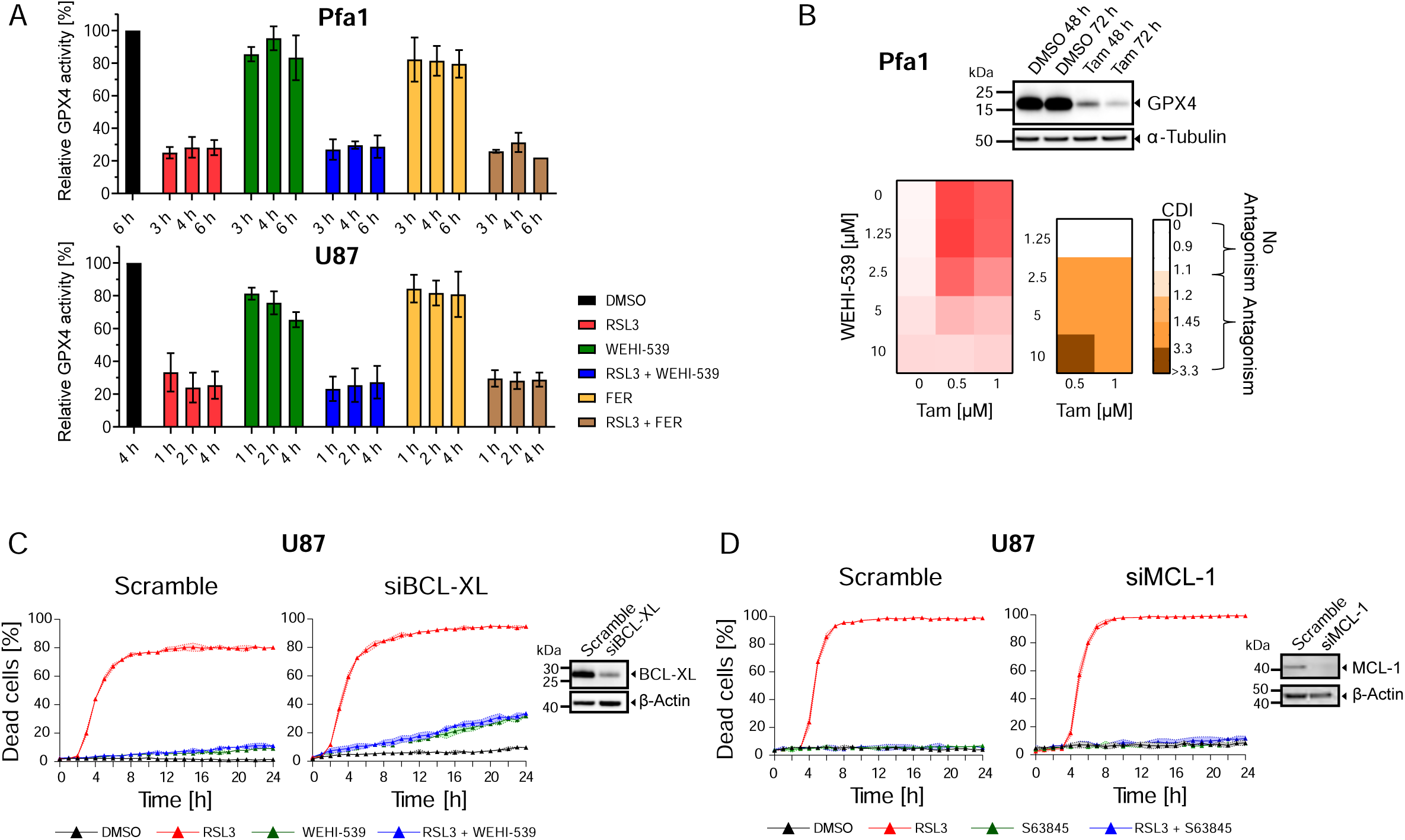
BH3 mimetics exhibit off-target activities that can suppress ferroptotic cell death. **(A)** Measurement of GPX4 activity. Cells were treated with 10 µM WEHI-539, 2 µM ferrostatin-1 (FER) and 100 nM RSL3 as indicated. Data show mean ± range of two technical replicates from one of two independent experiments. **(B)** Pfa1 cells with inducible GPX4 knock out were co-treated with WEHI-539 and tamoxifen (Tam) for 72 h. Cell death was determined by PI uptake. Antagonism was calculated by the Webb’s fractional product. GPX4 loss was confirmed by immunoblotting. Data show means from one representative out of three independent experiments. **(C)** U87 cells were treated with RSL3 (100 nM), WEHI-539 (10 μM), following depletion of BCL-XL expression. Cell death was determined by PI uptake performing live cell imaging. Data shown are means ± SD of three technical replicates from one out of two independent experiments. **(D)** U87 cells were treated with RSL3 (100 nM), S63845 (10 μM), following depletion of MCL-1 expression. Cell death was determined by PI uptake performing live cell imaging. Data shown are means ± SD of three technical replicates from one out of two independent experiments.

### BH3 mimetics suppress oxidative damage by intrinsic antioxidant activities

With the anti-ferroptotic activity of BH3-mimetics being off-target activities, it can be speculated that these compounds might have antioxidant activities similar to ferrostatin-1 and other radical trapping agents. To study the electrochemical properties of BH3-mimetics in comparison to ferrostatin-1, we employed differential pulse voltammetry measurements (**Fig.6A**). These measure the propensity of substances to accept or donate electrons under externally applied electric potentials. As such, voltammetry can describe the intrinsic redox activity of chemical compounds. Ferrostatin is known to easily donate electrons and prior voltammetry measurements in methanol environment have been reported before (Miotto et al., 2020). To compare ferrostatin-1 to BH3-mimetics, we here conducted these measurements in CH2Cl2 due to the otherwise limited solubility of BH3-mimetics at the concentrations required for comparative measurements. Ferrostatin-1 showed several, mostly overlapping oxidation peaks, with only the first peak being baseline separated. Such a propensity to donate electrons under positive electric potentials was seen for all BH3-mimetics, wherein WEHI-539 showed the closest resemblance to ferrostatin-1. AZD5991 and S63845 appeared unstable, as seen by noisy spectra and undefinable peaks at high positive voltages.

**Figure 6:**
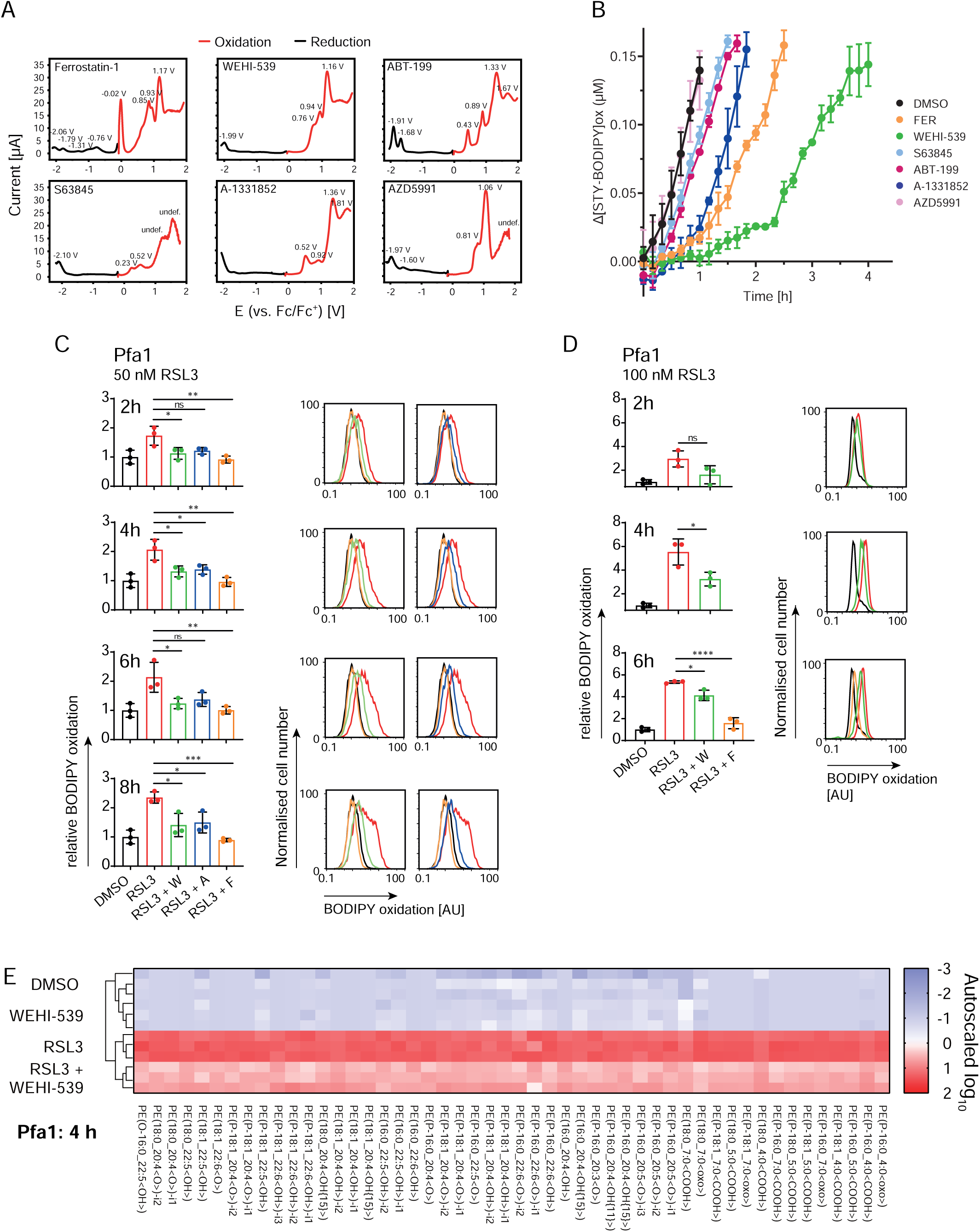
BH3 mimetics suppress oxidative damage by intrinsic antioxidant activities. **(A)** Voltammetry measurements for ferrostatin-1 (FER) and BH3-mimetics. Red and black curves correspond to the first oxidation or reduction measurement, respectively for each substrate. Peak undefined due to substrate deterioration (undef.). **(B)** FENIX assay to assess antioxidant activities of FER and BH3 mimetics by measuring STY-BODIPY (1 mM) co-autoxidation. All compounds were tested at 2 μM. Data were normalised to the DMSO control that lacked STY-BODIPY and are presented as the mean ± SD of three technical replicates from one out of three independent experiments. **C)** Pfa1 cells were stimulated with 50 nM RSL3, 10 µM WEHI-539 (W),10 µM A-1331852 (A) and 2 µM ferrostatin-1 (F) for the indicated times before they were analysed for lipid peroxidation (C11 BODIPY 581/591 conversion) via flow cytometry. Data shown are mean ± SD of three independent experiments. One-way ANOVA with Tukey‘s multiple comparisons test * = p ≤ 0.05, **** = p ≤ 0.0001, ns = not significant. Overlay graphs are from one representative experiment. DMSO control (0h) is the same in all bar graphs and plots. **(E)** Lipid oxidation measurements by LC-MS/MS. Pfa1 cells were treated with 10 µM WEHI-539 and 100 nM RSL3 alone or in combination for 4 h. Normalised peak areas of oxidised lipids showing significant regulation (ANOVA, adjusted p-value [FDR] cutoff: 0.05) were log transformed and visualised using heatmaps. The color scheme corresponds to auto-scaled log fold change relative to the mean log value within the samples. PE, phosphatidylethanolamines, i indicates isomeric lipids. Data are from one experiment with three technical replicates.

As *in vitro* readouts for the potency of BH3-mimetics to interfere with self-propagating lipid peroxidation, we compared these to ferrostatin-1 in FENIX assays. Tested at equimolar concentrations, especially the BCL-XL inhibitors WEHI-539 and A-1331852 showed substantial antioxidant activities, whereas AZD5991 was the only BH3-mimetic not separable from the negative control (**Fig.6B**). Kinetic analyses *in cellulo* demonstrated that both WEHI-539 and A-1331852 effectively suppressed BODIPY oxidation in Pfa1 cells when treated with 50 nM RSL3 (**Fig.6C, Supplemental Fig.9A**). Similar results were obtained in U87 cells, yet A-1331852 was less effective (**Supplemental Fig.9B,C**). At higher concentrations of RSL3 (100 nM), BODIPY oxidation gradually proceeded in the presence of WEHI-539 to become indistinguishable from the positive control after 4-6 h in Pfa1 cells (**Fig.6D**). However, despite full BODIPY conversion within this period, which would widely be interpreted as a surrogate indicator for maximal lipid peroxidation, RSL3/WEHI-539 combination treatments clearly did not kill these cells within 24 h of treatment (**Fig.3C**). In contrast to the accumulative oxidation of a fixed amount of dye over time, the rate of lipid turnover in living cells might instead permit RSL3/WEHI-539 treated cells to limit the abundance of oxidised lipids to sublethal amounts that do not compromise membrane integrity. We therefore also performed direct lipidomic analyses, which indeed demonstrated that the overall extent of RSL3-induced lipid oxidation was substantially reduced in presence of WEHI-539 (**Fig.6E**).

Overall, these data demonstrate that most BH3-mimetics possess intrinsic anti-oxidant activities, with A-1331852 and WEHI-539 showing the strongest effect. Furthermore, these antioxidant activities suffice to reduce lipid oxidation in living cells substantially, affording the survival of sublethally damaged cells.

## Discussion

Here we demonstrate that ferroptosis and apoptosis can manifest as coupled cell death modalities, and furthermore that cell death sensitisation by BH3-mimetics can redirect cell fate outcomes from ferroptosis to apoptosis. Surprisingly, previously undescribed antioxidant properties of BH3-mimetics can counter-intuitively suppress ferroptosis in cells that do not rely on respective BH3-mimetic targets for apoptosis prevention and cell survival under ferroptotic stress.

Hallmarks of apoptosis, such as cytochrome-c release into the cytosol, effector caspase activation and (transient) membrane blebbing, can be observed in cells exposed to ferroptotic stress. This might have remained obscure so far due to prior studies often lacking sufficiently detailed kinetic studies of cell death manifestation under inclusion of appropriate control conditions, or due to applying extreme ferroptotic stress conditions that molecularly and phenotypically left no space for apoptotic hallmarks to manifest prominently. As we show, apoptosis only partially contributed to overall cell death in HT1080 cells, the most widely used model system to study ferroptosis. The cellular phenotype of apoptotic membrane blebbing (Kerr et al., 1972) often was observed only transiently, before being overridden by cell inflation and membrane rupture. The latter is also observed during apoptosis as a delayed “secondary necrosis” phenotype, especially in absence of the clearing of cellular debris by neighbouring cells or macrophages (Silva, 2010). However, under ferroptotic stress the features of membrane blebbing and cell lysis kinetically appear tightly coupled and indeed can co-occur in time. Ferroptotic membrane damage therefore likely destabilises membranes to such an extent that apoptotic blebs might lose integrity while cells have already entered the execution phase of apoptosis, thus resulting in rapid cell lysis. Cell lysis is also the endpoint of necroptosis and pyroptosis, two forms of cell death which exhibit strong interplay with apoptosis and during which protein pores permeabilise the plasma membrane (Tsuchiya et al., 2019, Liu et al., 2016, Newton et al., 2024, Yuan and Ofengeim, 2024). Membrane damage by ferroptotic lipid oxidation might promote such lytic cell death modalities in combinatorial stress conditions, as shown for example for gasdermin D-mediated pyroptosis (Kang et al., 2018), especially if lipid oxidation eases the insertion and interaction of pore forming proteins into and within the plasma membrane. This equally might apply for BAX/BAK-dependent pore formation as a key event during apoptosis signalling at mitochondrial membranes. Unsaturated lipids support BAX and BAK pore formation in membranes (Dadsena et al., 2024), and ferroptotic oxidative damage to such lipids might further support the formation of BAX/BAK pores. Indeed, we show here that the presence of BAX/BAK is crucial for apoptosis engagement under ferroptotic oxidative stress. Radical stress is indeed long known to promote apoptosis (Korsmeyer et al., 1995). A direct mitochondrial link between GPX4 activity and apoptosis resistance under radical stress was also suggested, where GPX4 suppresses the peroxidation of the mitochondrial lipid cardiolipin and prevents the release of cytochrome-c (Nomura et al., 2000), yet these findings were not validated independently so far.

As shown here, RSL3-induced ferroptotic stress suffices to release submaximal amounts of cytochrome-c from mitochondria into the cytosol, and therefore contrasts with canonical apoptosis induction, where BH3-only proteins induce cytochrome-c release that proceeds in an all-or-none manner and which triggers highly effective effector caspase activation (Goldstein et al., 2000, Rehm et al., 2006). Limited cytochrome-c release, as seen in our study, in many cases is sufficient to trigger the manifestation of apoptotic hallmarks. This is in contrast to scenarios of residual cytochrome-c release upon low or transient apoptotic stress that nevertheless causes sublethal damage due to effector caspase activation (Ichim et al., 2015, Gradzka-Boberda et al., 2022). Still, the submaximal cytochrome-c release observed upon RSL3-induced ferroptotic stress likely explains why apoptosis cannot be observed as the primary but nevertheless as a significant contributing cell death modality in our experimental conditions.

To provoke BAX/BAK-dependent permeabilisation of the outer mitochondrial membrane, BH3-mimetics are widely used research tools as well as therapeutically relevant agents (Czabotar and Garcia-Saez, 2023, Diepstraten et al., 2022). We identified that both synergistic interactions with GPX4 inhibition and surprisingly also the opposite, prevention of cell death, could be observed. It is conceivable that lipid oxidation, which as explained above could enhance apoptosis sensitivity, induces a stronger reliance on cell line-specific subsets of anti-apoptotic BCL-2 family members for survival, and that antagonising these family members results in synergistic rerouting of cell death towards apoptosis. This is in line with a previous study that reported the sensitisation of acute myeloid leukemia cells to ABT-199/venetoxclax upon exposure to RSL3 (Lin et al., 2019). Where BH3-mimetics are applied that do not target anti-apoptotic family members required to withstand radical stress, anti-oxidative properties of such BH3-mimetics likely prevail and suppress cell death, as evidenced here by reduced C11 BODIPY and lipid oxidation in cells surviving GPX4 inhibition. Antioxidant properties that reduce cellular ferroptosis susceptibility have been described for various bioactive compounds in a large screening effort in HT1080 cells before (Conlon et al., 2021), yet BH3-mimetics were not studied systematically and across different cell lines. Furthermore, since combination studies with BH3-mimetics and ferroptosis stressors were mostly limited to BCL-2 antagonism (Lin et al., 2019, Yu et al., 2024), and since ABT-199/venetoclax exhibits the least antioxidant activity, the ferroptosis suppressing action of BH3-mimetics has been missed so far. Lipid oxidation upon ferroptotic stress as well as the steady-state lipid oxidation status are not commonly measured directly, and instead most often are determined by the oxidation of C11 BODIPY 581/591 as a surrogate fluorescence marker. While suitable for determining the overall, accumulated lipid oxidation, endpoint BODIPY measurements might not reflect the actual lipid oxidation status in cells surviving ferroptotic stress. As lipid turnover and membrane repair continuously counteract membrane damage (Barisch et al., 2023, Mathiowetz and Olzmann, 2024), high BODIPY signals, especially after prolonged observation times, might overestimate actual lipid oxidation status. Indeed, we observed cells with high endpoint amounts BODIPY oxidation to survive treatment, indicating the requirement for direct lipidomics measurements to reliably assess oxidative lipid damage.

To conclude, likely dependent on the composition of the BCL-2 family interactome and cell line-specific intrinsic apoptotic priming (Ni Chonghaile et al., 2011), synergistically enhanced cell death or unexpected cell survival in presence of BH3-mimetics and ferroptotic stress will be the predominant outcome. Our work therefore highlights the need to assess case-specifically if combinations of ferroptosis inducers and BH3-mimetics can be suitable combinations to enhance cancer cell killing or if these will rather result in unexpected and unwanted cell survival. This is of particular relevance since we found that cells maintain or regain proliferative capacity in presence of BH3-mimetics with pronounced antioxidant activities.

## Acknowledgements

YQ received support from the China Scholarship Council (No. 202008320310). MR receives funding from the Deutsche Forschungsgemeinschaft (DFG) under Germany’s Excellence Strategy (EXC 2075, project number 390740016) and through DFG grant MO 3226/4-1. MR, AG and MC furthermore receive support through DFG grant TRR353/1 – 471011418 (Regulation of cell death decisions), and MC from the European Research Council (ERC) under the European Union’s Horizon 2020 research and innovation programme (grant agreement No. GA 884754).

The authors thank Prof Dr Sabine Ludwigs (Institute of Polymer Chemistry, University of Stuttgart) for providing access to the differential pulse voltammetry equipment, as well as Dennis Großmann and Svenja Bechthold for general support throughout the voltammetry measurements. The authors also acknowledge the Technology Platform “Cellular Analytics“ of the Stuttgart Research Center Systems Biology for their support & assistance.

## Author contributions

YQ, data acquisition, analysis and interpretation, draft writing; JH, data acquisition, analysis and interpretation, draft writing; BW, data acquisition, analysis and interpretation, draft writing; AR, data acquisition, analysis, draft revision; RT, data acquisition, analysis, draft revision; AG, data interpretation, draft revision; SL, data interpretation, supervision, draft revision; MC, data interpretation, supervision, draft revision; DS, data acquisition, analysis and interpretation, supervision, conception and design, draft writing and revision; MR, data interpretation, draft writing, supervision, funding acquisition, draft writing and revision, overall conception and design of the work.

## Supplemental Movie 1

HT1080S cells were stimulated with 2.25 µM RSL3 and monitored over a time period of 24 h. Images were taken every 10 min at 20x magnification. Medium contained PI. One representative movie of three independent experiments is shown.

---

**Supplemental Figure 1.**
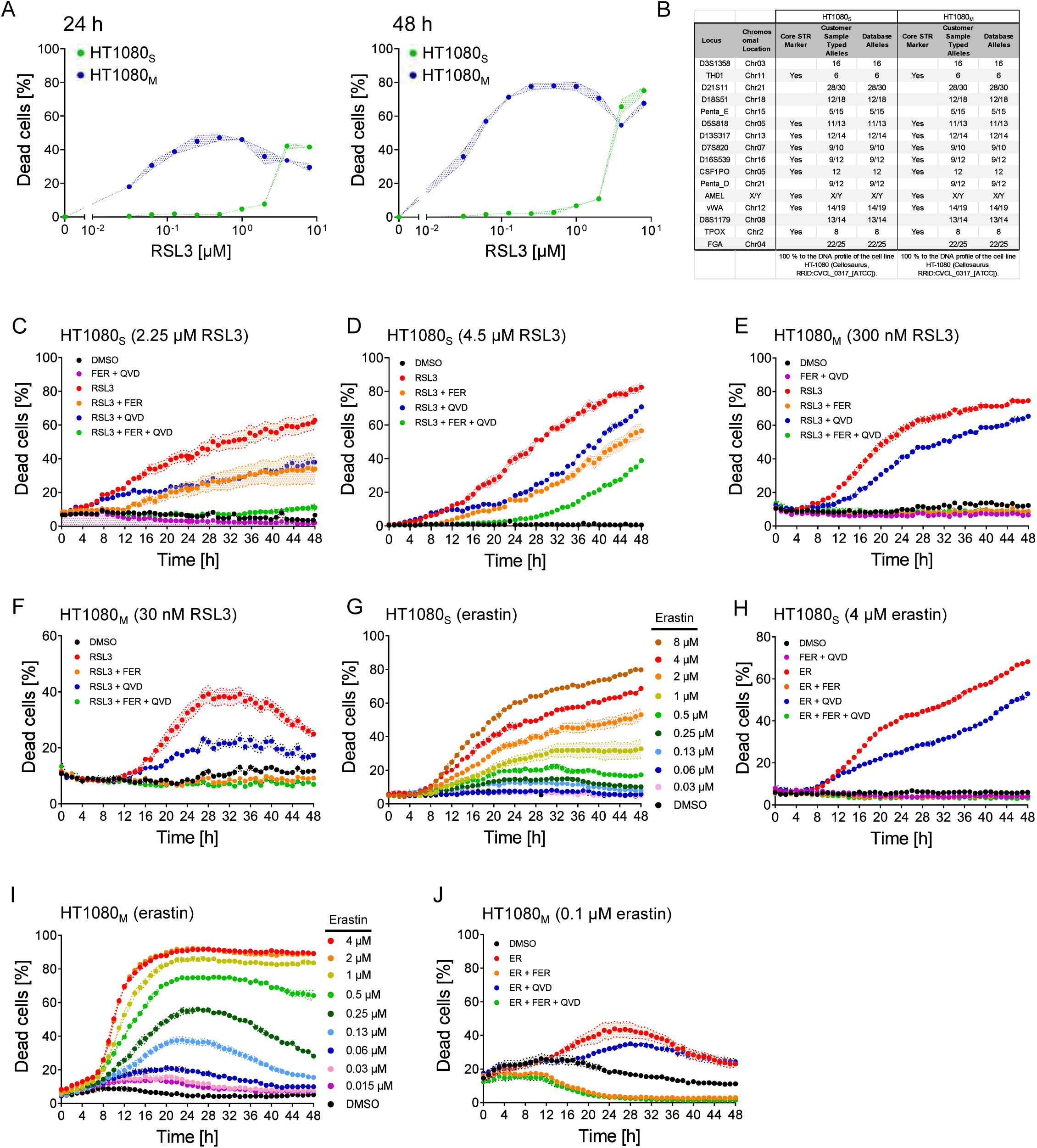
(A) Quantification of cell death, calculated as percentage of PI-positive cells. Data show mean ± range of technical duplicates of one out of two independent experiments. (B) Profiling of HT1080S and HT1080M strains using highly polymorphic short tandem repeat loci (STRs) analysis. **(C-J)** Quantification of cell death as in **(A)**. cells. Cells were stimulated with the indicated concentrations of RSL3 and erastin as well as with 50 µM QVD and 2 µM ferrostatin-1 (FER). Data are means ± range of technical duplicates **(D)** and mean ± SEM of technical triplicates (rest) from one out of three independent experiments.

**Supplemental Figure 2.**
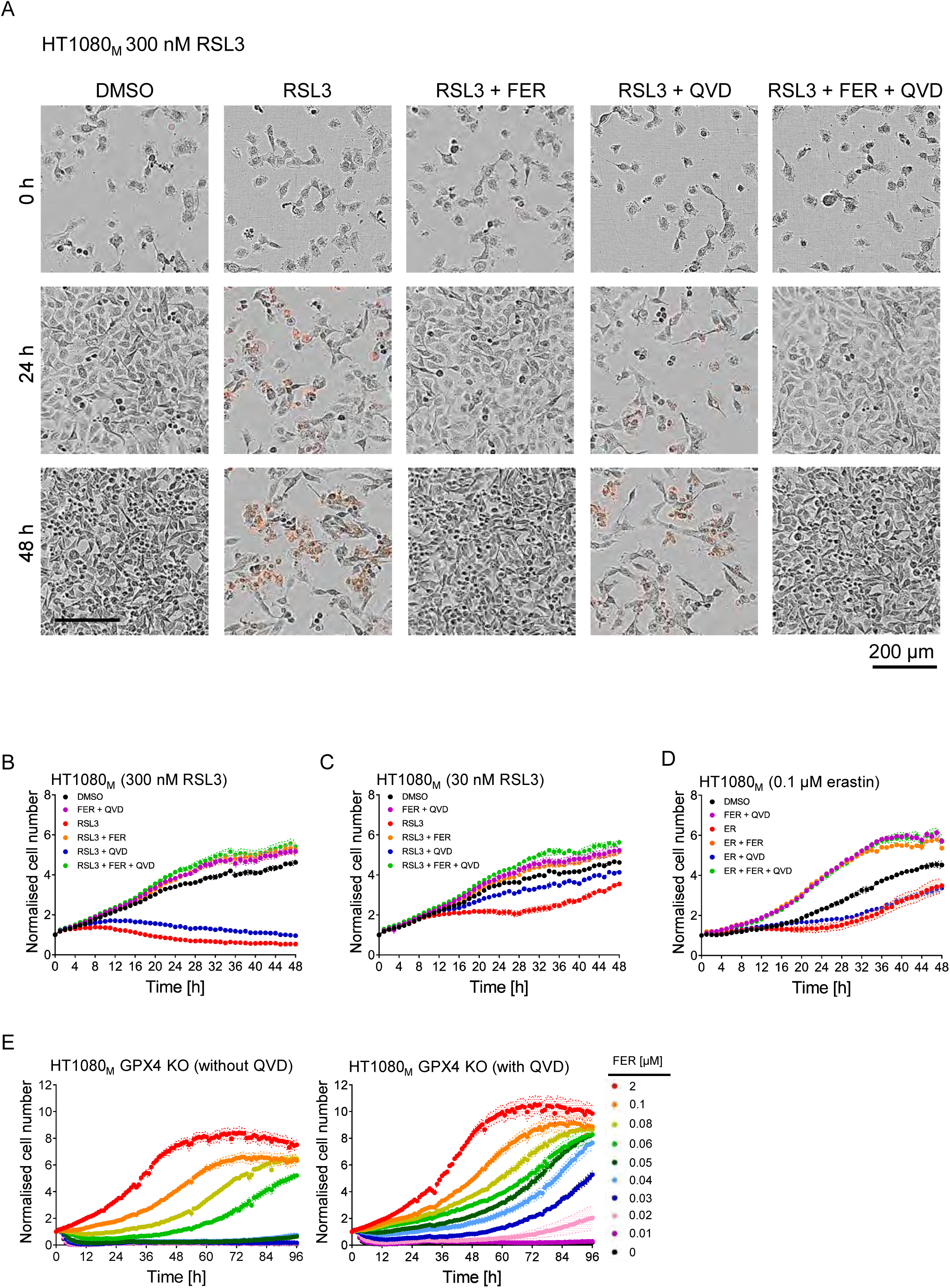
**(A)** Images were taken at 10x magnification and are from one representative of three independent experiments. Panels show brightfield overlays with PI fluorescence. **(B-D)** Cell proliferation. Cells were stimulated with the indicated concentrations of RSL3 and erastin as well as with 50 µM QVD and 2 µM ferrostatin-1 (FER). Data are means ± SEM of technical triplicates from one out of three independent experiments. **(E)** Cell proliferation was determined in HT1080M cells lacking GPX4. QVD was used at 50 µM. Data show means ± SEM of technical triplicates from one out of three independent experiments.

**Supplemental Figure 3.**
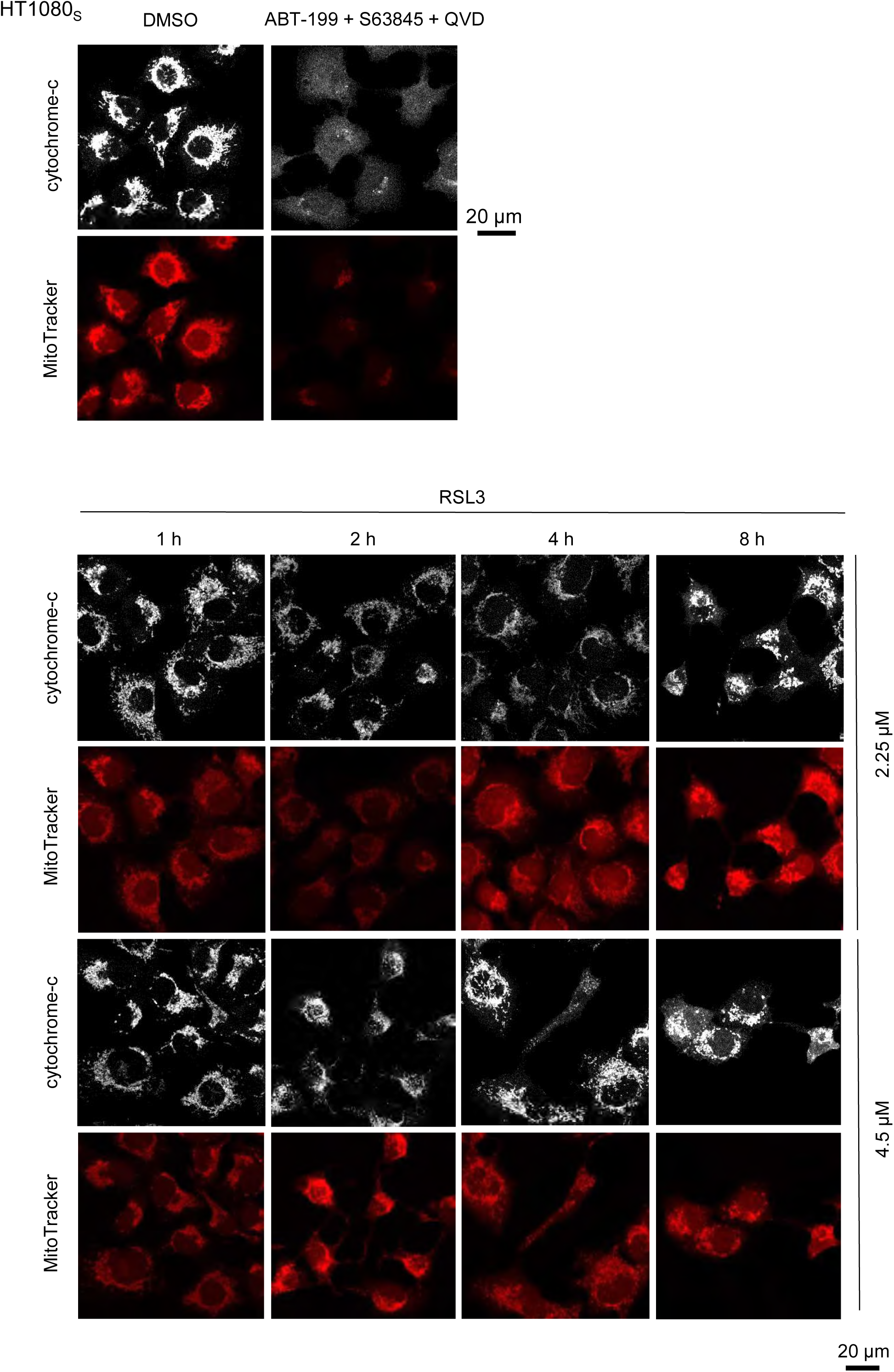
Cells loaded with MitoTracker Red CMXRos were stimulated with the indicated concentrations of RSL3 and with 10 µM ABT-199, 10 µM S63845 and 50 µM QVD for 4 h as an apoptosis positive control. At the respective time points, cells were fixed and immunostained for cytochrome-c. Images are from one representative of three independent experiments.

**Supplemental Figure 4.**
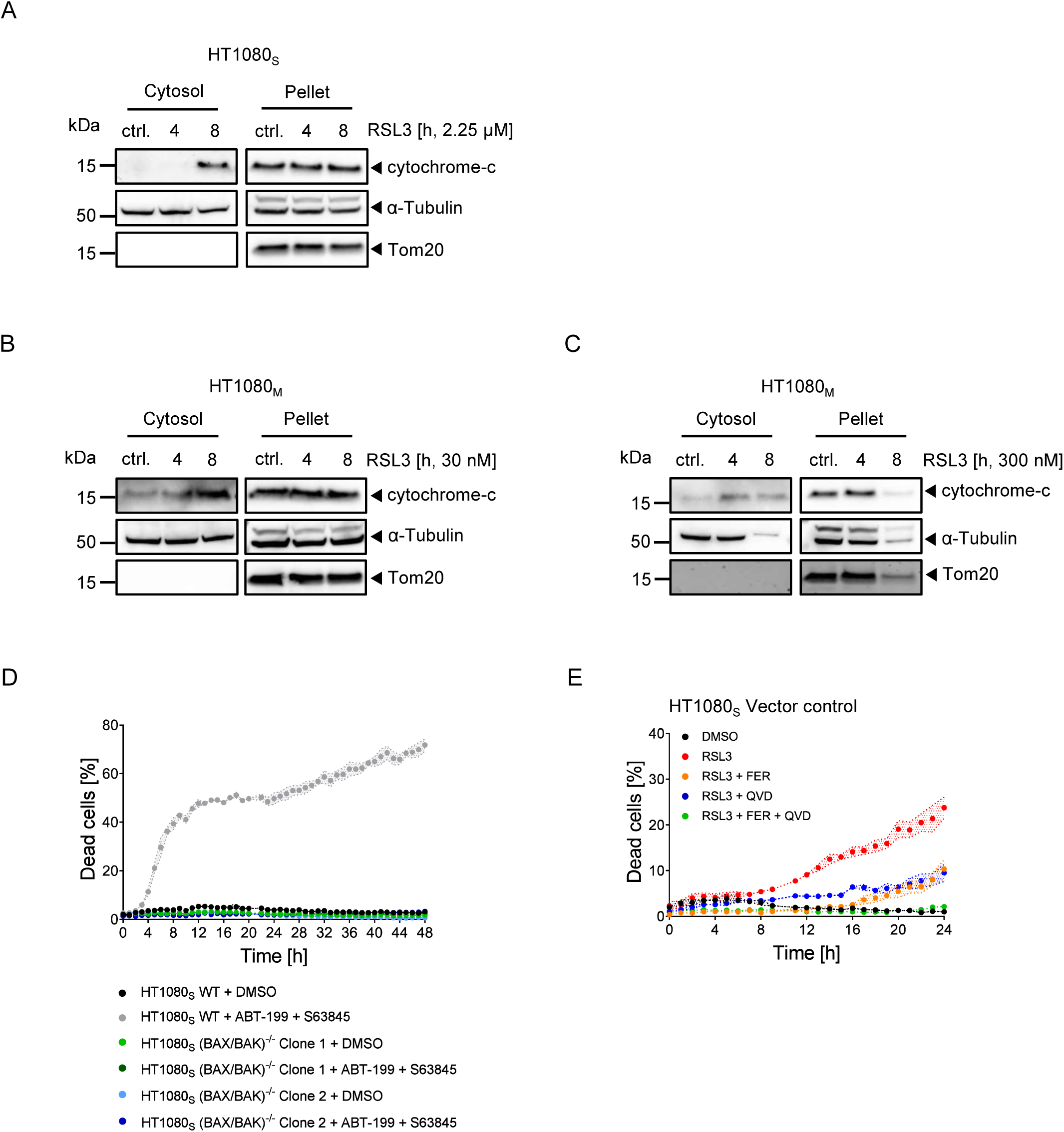
(A-C) Cells were fractioned into cytoplasm and pellet containing the mitochondria. Fractions were blotted for cytochrome-c. One representative of three (HT1080S) or two (HT1080M) independent experiments is shown. **(D)** Quantification of cell death, calculated as percentage of PI-positive cells. Cells were stimulated with 10 µM ABT-199 + 10 µM S63845. Data are means ± range of technical duplicates from one out of three independent experiments. **(E)** Quantification of cell death, calculated as percentage of PI-positive cells. Cells were stimulated with 2.25 µM RSL3, 50 µM QVD and 2 µM ferrostatin-1 (FER). Data are means ± range of technical duplicates from one out of four independent experiments.

**Supplemental Figure 5.**
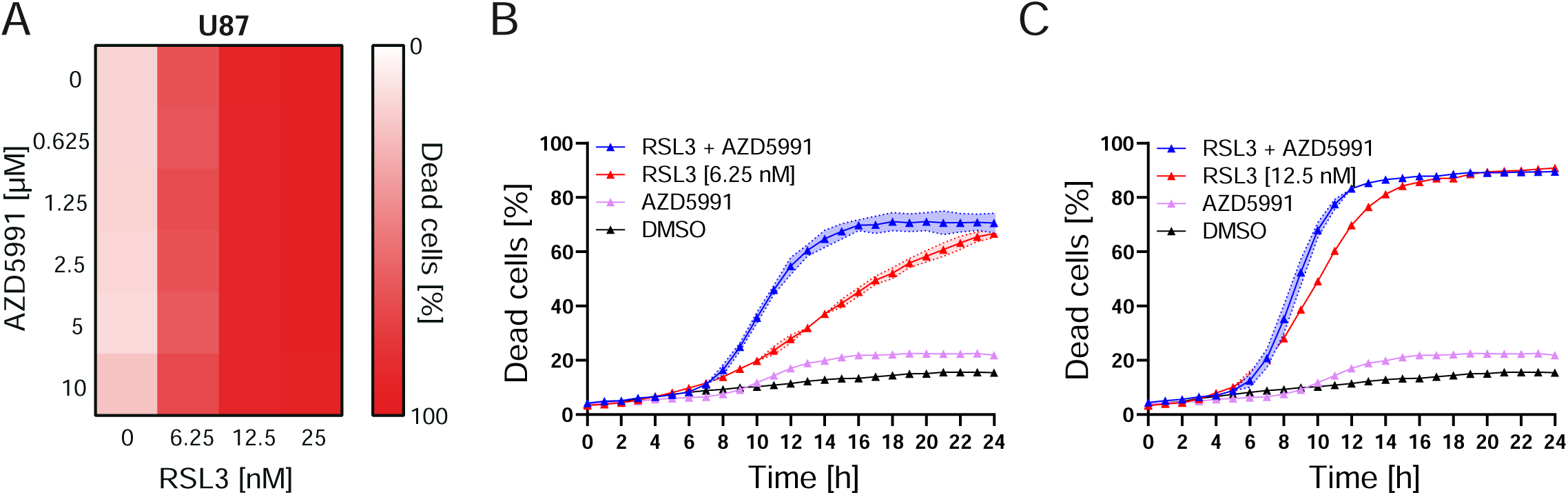
**(A)** U87 cells were treated as indicated. Cell death was determined by the uptake of PI at 24 h. Heatmap shows the mean of three independent experiments. **(B, C)** Cells were treated with RSL3 and AZD5991 (10 µM). Cell death was determined by PI uptake using time-lapse imaging. Data show means ± SD from three independent experiments

**Supplemental Figure 6.**
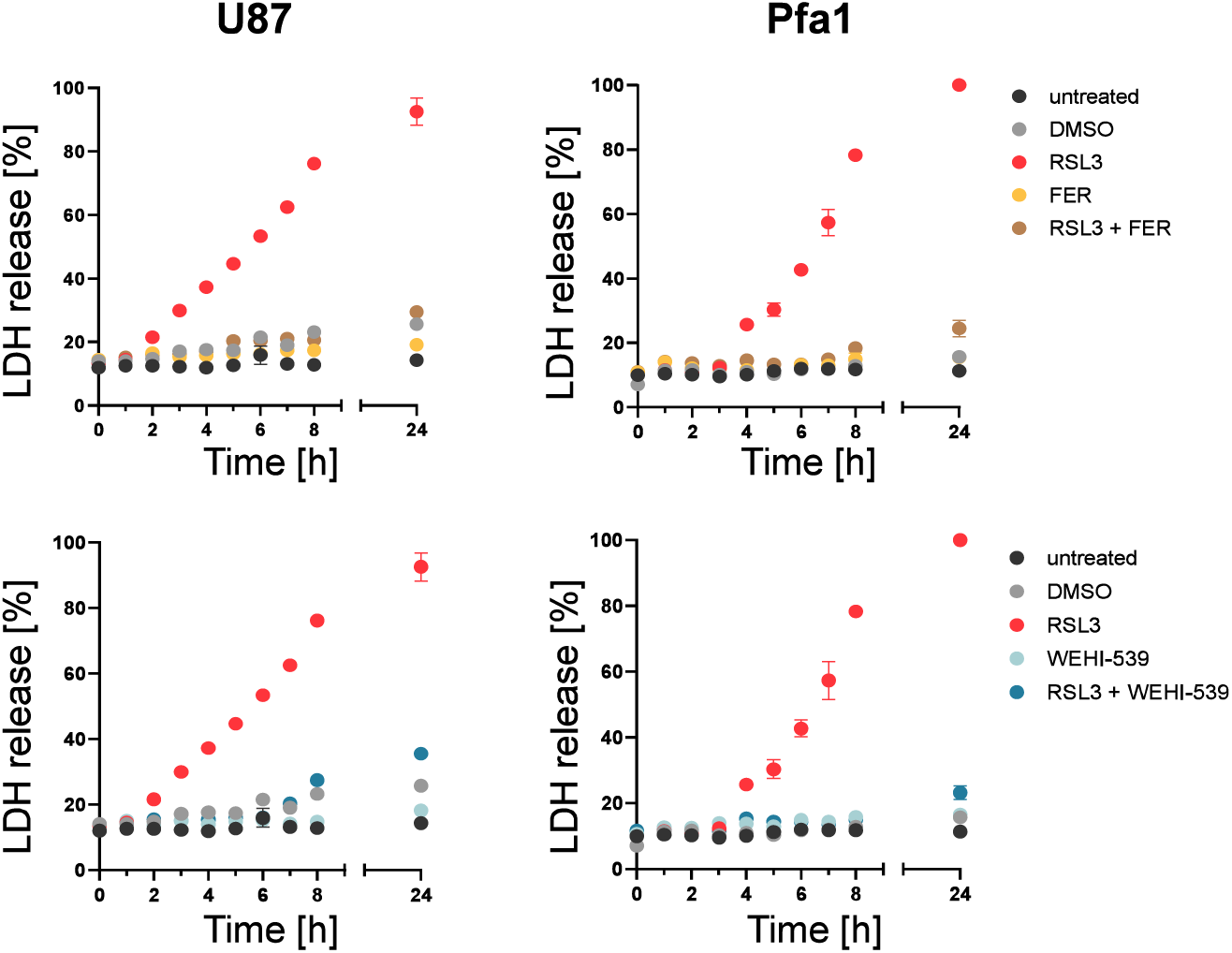
LDH release was measured following treatments with DMSO, 10 µM WEHI-539, 2 µM ferrostatin-1 (FER), or 100 nM of RSL3 alone or in combination. Data are means ± range from one experiment.

**Supplemental Figure 7.**
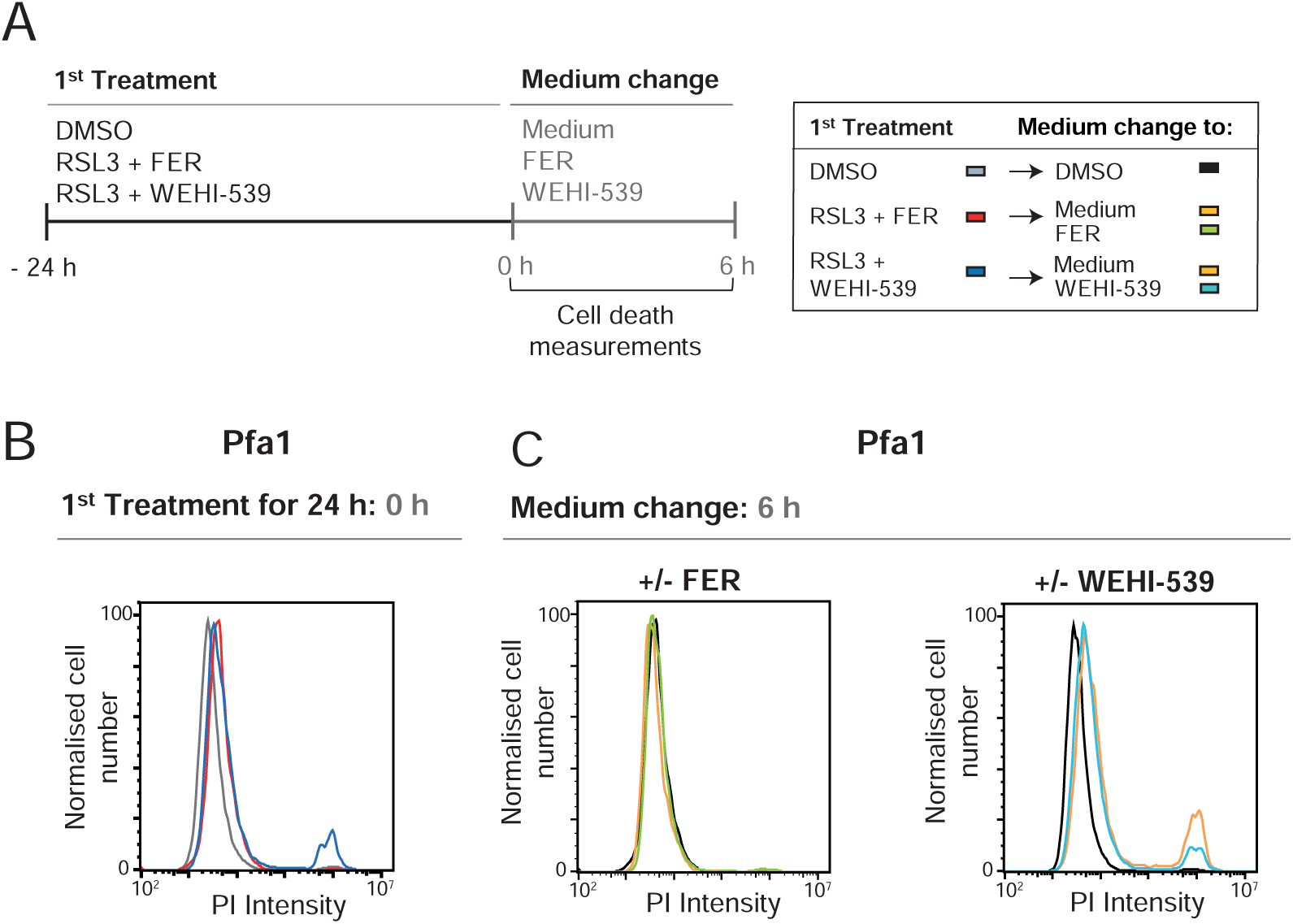
**(A)** Treatment schedule. **(B, C)** Pfa1 cells received a 1st treatment with DMSO or a combination treatment of RSL3 (100 nM) with WEHI-539 (10 µM) or RSL3 with ferrostatin-1 (FER, 2 µM) for 24 h. Cells were then washed with PBS and the medium was changed as indicated in (A). Cell death was measured by PI uptake after the first treatment and 6 h after the medium change. One representative experiment of three independent experiments is shown.

**Supplemental Figure 8.**
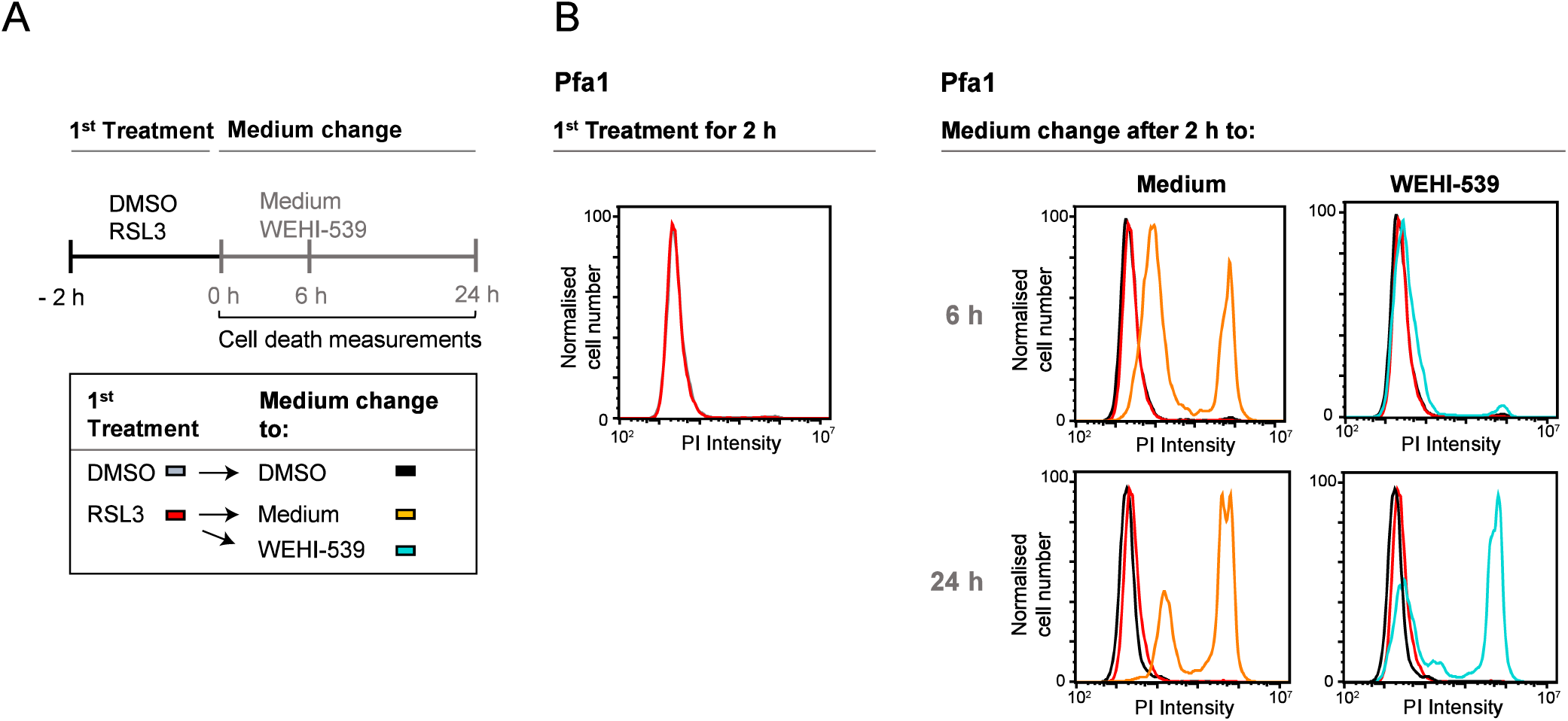
**(A)** Treatment schedule **(B, C)** Cells received the 1st treatment with DMSO or RSL3 (100 nM) for 2 h. Cells were then washed with PBS and the medium was changed as indicated. Cell death was measured by PI uptake at 2 h (B) as well as 6 h and 24 h after the medium change. One out of two independent experiments is shown.

**Supplemental Figure 9.**
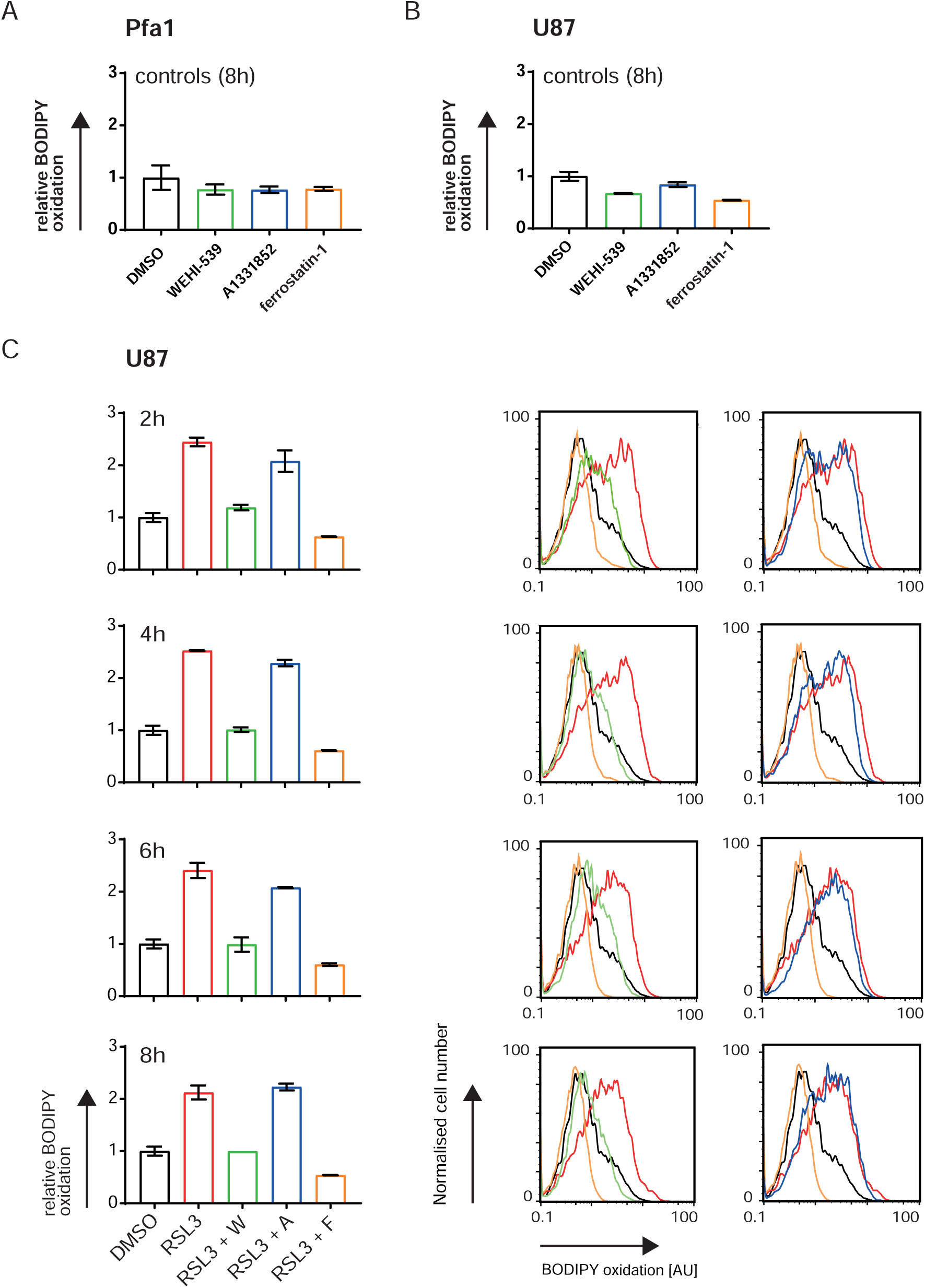
**A)** Cells were stimulated with 50 nM RSL3, 10 µM WEHI-539, 10 µM A-1331852 and 2 µM ferrostatin-1. Shown are the controls, mean ± SD of three independent experiments. **B, C)** U87 cells were stimulated with 12.5 nM RSL3, 10 µM WEHI-539 (W), 10 µM A-1331852 (A) and 2 µM ferrostatin-1 (F) for the indicated times. Data shown are mean ± range of two independent experiments. Overlay graphs are from one representative experiment. DMSO control (8h) is the same in all bar graphs and plots.

## References

Barisch, C., Holthuis, J. C. M. & Cosentino, K. 2023. Membrane damage and repair: a thin line between life and death. Biol Chem, 404, 467–490.

Chu, J., Liu, C. X., Song, R. & Li, Q. L. 2020. Ferrostatin-1 protects HT-22 cells from oxidative toxicity. Neural Regen Res, 15, 528–536.

Conlon, M., Poltorack, C. D., Forcina, G. C., Armenta, D. A., Mallais, M., Perez, M. A., Wells, A., Kahanu, A., Magtanong, L., Watts, J. L., Pratt, D. A. & Dixon, S. J. 2021. A compendium of kinetic modulatory profiles identifies ferroptosis regulators. Nat Chem Biol, 17, 665–674.

Czabotar, P. E. & Garcia-Saez, A. J. 2023. Mechanisms of BCL-2 family proteins in mitochondrial apoptosis. Nat Rev Mol Cell Biol, 24, 732–748.

Dadsena, S., Cuevas Arenas, R., Vieira, G., Brodesser, S., Melo, M. N. & Garcia-Saez, A. J. 2024. Lipid unsaturation promotes BAX and BAK pore activity during apoptosis. Nat Commun, 15, 4700.

Diepstraten, S. T., Anderson, M. A., Czabotar, P. E., Lessene, G., Strasser, A. & Kelly, G. L. 2022. The manipulation of apoptosis for cancer therapy using BH3-mimetic drugs. Nat Rev Cancer, 22, 45–64.

Dixon, S. J., Patel, D. N., Welsch, M., Skouta, R., Lee, E. D., Hayano, M., Thomas, A. G., Gleason, C. E., Tatonetti, N. P., Slusher, B. S. & Stockwell, B. R. 2014. Pharmacological inhibition of cystine-glutamate exchange induces endoplasmic reticulum stress and ferroptosis. Elife, 3, e02523.

Doll, S., Freitas, F. P., Shah, R., Aldrovandi, M., Da Silva, M. C., Ingold, I., Goya Grocin, A., Xavier Da Silva, T. N., Panzilius, E., Scheel, C. H., Mourao, A., Buday, K., Sato, M., Wanninger, J., Vignane, T., Mohana, V., Rehberg, M., Flatley, A., Schepers, A., Kurz, A., White, D., Sauer, M., Sattler, M., Tate, E. W., Schmitz, W., Schulze, A., O’donnell, V., Proneth, B., Popowicz, G. M., Pratt, D. A., Angeli, J. P. F. & Conrad, M. 2019. FSP1 is a glutathione-independent ferroptosis suppressor. Nature, 575, 693–698.

Dolma, S., Lessnick, S. L., Hahn, W. C. & Stockwell, B. R. 2003. Identification of genotype-selective antitumor agents using synthetic lethal chemical screening in engineered human tumor cells. Cancer Cell, 3, 285–96.

Dondelinger, Y., Priem, D., Huyghe, J., Delanghe, T., Vandenabeele, P. & Bertrand, M. J. M. 2023. NINJ1 is activated by cell swelling to regulate plasma membrane permeabilization during regulated necrosis. Cell Death Dis, 14, 755.

Eagle, H. 1955. The specific amino acid requirements of a mammalian cell (strain L) in tissue culture. J Biol Chem, 214, 839–52.

Goldstein, J. C., Waterhouse, N. J., Juin, P., Evan, G. I. & Green, D. R. 2000. The coordinate release of cytochrome c during apoptosis is rapid, complete and kinetically invariant. Nat Cell Biol, 2, 156–62.

Gradzka-Boberda, S., Gentle, I. E. & Hacker, G. 2022. Pattern Recognition Receptors of Nucleic Acids Can Cause Sublethal Activation of the Mitochondrial Apoptosis Pathway during Viral Infection. J Virol, 96, e0121222.

Hirata, Y., Cai, R., Volchuk, A., Steinberg, B. E., Saito, Y., Matsuzawa, A., Grinstein, S. & Freeman, S. A. 2023. Lipid peroxidation increases membrane tension, Piezo1 gating, and cation permeability to execute ferroptosis. Curr Biol, 33, 1282–1294 e5.

Ichim, G., Lopez, J., Ahmed, S. U., Muthalagu, N., Giampazolias, E., Delgado, M. E., Haller, M., Riley, J. S., Mason, S. M., Athineos, D., Parsons, M. J., Van De Kooij, B., Bouchier-Hayes, L., Chalmers, A. J., Rooswinkel, R. W., Oberst, A., Blyth, K., Rehm, M., Murphy, D. J. & Tait, S. W. 2015. Limited mitochondrial permeabilization causes DNA damage and genomic instability in the absence of cell death. Mol Cell, 57, 860–72.

Kang, R., Zeng, L., Zhu, S., Xie, Y., Liu, J., Wen, Q., Cao, L., Xie, M., Ran, Q., Kroemer, G., Wang, H., Billiar, T. R., Jiang, J. & Tang, D. 2018. Lipid Peroxidation Drives Gasdermin D-Mediated Pyroptosis in Lethal Polymicrobial Sepsis. Cell Host Microbe, 24, 97–108 e4.

Kerr, J. F., Wyllie, A. H. & Currie, A. R. 1972. Apoptosis: a basic biological phenomenon with wide-ranging implications in tissue kinetics. Br J Cancer, 26, 239–57.

Korsmeyer, S. J., Yin, X. M., Oltvai, Z. N., Veis-Novack, D. J. & Linette, G. P. 1995. Reactive oxygen species and the regulation of cell death by the Bcl-2 gene family. Biochim Biophys Acta, 1271, 63–6.

Lin, K. H., Xie, A., Rutter, J. C., Ahn, Y. R., Lloyd-Cowden, J. M., Nichols, A. G., Soderquist, R. S., Koves, T. R., Muoio, D. M., Maciver, N. J., Lamba, J. K., Pardee, T. S., Mccall, C. M., Rizzieri, D. A. & Wood, K. C. 2019. Systematic Dissection of the Metabolic-Apoptotic Interface in AML Reveals Heme Biosynthesis to Be a Regulator of Drug Sensitivity. Cell Metab, 29, 1217–1231 e7.

Liu, X., Zhang, Z., Ruan, J., Pan, Y., Magupalli, V. G., Wu, H. & Lieberman, J. 2016. Inflammasome-activated gasdermin D causes pyroptosis by forming membrane pores. Nature, 535, 153–8.

Lu, S. C. 2013. Glutathione synthesis. Biochim Biophys Acta, 1830, 3143–53.

Mandal, P. K., Seiler, A., Perisic, T., Kolle, P., Banjac Canak, A., Forster, H., Weiss, N., Kremmer, E., Lieberman, M. W., Bannai, S., Kuhlencordt, P., Sato, H., Bornkamm, G. W. & Conrad, M. 2010. System x(c)- and thioredoxin reductase 1 cooperatively rescue glutathione deficiency. J Biol Chem, 285, 22244–53.

Mathiowetz, A. J. & Olzmann, J. A. 2024. Lipid droplets and cellular lipid flux. Nat Cell Biol, 26, 331–345.

Matyash, V., Liebisch, G., Kurzchalia, T. V., Shevchenko, A. & Schwudke, D. 2008. Lipid extraction by methyl-tert-butyl ether for high-throughput lipidomics. J Lipid Res, 49, 1137–46.

Miotto, G., Rossetto, M., Di Paolo, M. L., Orian, L., Venerando, R., Roveri, A., Vuckovic, A. M., Bosello Travain, V., Zaccarin, M., Zennaro, L., Maiorino, M., Toppo, S., Ursini, F. & Cozza, G. 2020. Insight into the mechanism of ferroptosis inhibition by ferrostatin-1. Redox Biol, 28, 101328.

Mishima, E., Ito, J., Wu, Z., Nakamura, T., Wahida, A., Doll, S., Tonnus, W., Nepachalovich, P., Eggenhofer, E., Aldrovandi, M., Henkelmann, B., Yamada, K. I., Wanninger, J., Zilka, O., Sato, E., Feederle, R., Hass, D., Maida, A., Mourao, A. S. D., Linkermann, A., Geissler, E. K., Nakagawa, K., Abe, T., Fedorova, M., Proneth, B., Pratt, D. A. & Conrad, M. 2022. A non-canonical vitamin K cycle is a potent ferroptosis suppressor. Nature, 608, 778–783.

Neitemeier, S., Jelinek, A., Laino, V., Hoffmann, L., Eisenbach, I., Eying, R., Ganjam, G. K., Dolga, A. M., Oppermann, S. & Culmsee, C. 2017. BID links ferroptosis to mitochondrial cell death pathways. Redox Biol, 12, 558–570.

Newton, K., Strasser, A., Kayagaki, N. & Dixit, V. M. 2024. Cell death. Cell, 187, 235–256.

Ni Chonghaile, T., Sarosiek, K. A., Vo, T. T., Ryan, J. A., Tammareddi, A., Moore Vdel, G., Deng, J., Anderson, K. C., Richardson, P., Tai, Y. T., Mitsiades, C. S., Matulonis, U. A., Drapkin, R., Stone, R., Deangelo, D. J., Mcconkey, D. J., Sallan, S. E., Silverman, L., Hirsch, M. S., Carrasco, D. R. & Letai, A. 2011. Pretreatment mitochondrial priming correlates with clinical response to cytotoxic chemotherapy. Science, 334, 1129–33.

Nomura, K., Imai, H., Koumura, T., Kobayashi, T. & Nakagawa, Y. 2000. Mitochondrial phospholipid hydroperoxide glutathione peroxidase inhibits the release of cytochrome c from mitochondria by suppressing the peroxidation of cardiolipin in hypoglycaemia-induced apoptosis. Biochem J, 351, 183–93.

Redza-Dutordoir, M. & Averill-Bates, D. A. 2016. Activation of apoptosis signalling pathways by reactive oxygen species. Biochim Biophys Acta, 1863, 2977–2992.

Rehm, M., Huber, H. J., Dussmann, H. & Prehn, J. H. 2006. Systems analysis of effector caspase activation and its control by X-Linked inhibitor of apoptosis protein. Embo J, 25, 4338–49.

Seiler, A., Schneider, M., Forster, H., Roth, S., Wirth, E. K., Culmsee, C., Plesnila, N., Kremmer, E., Radmark, O., Wurst, W., Bornkamm, G. W., Schweizer, U. & Conrad, M. 2008. Glutathione peroxidase 4 senses and translates oxidative stress into 12/15-lipoxygenase dependent- and Aif-Mediated cell death. Cell Metab, 8, 237–48.

Silva, M. T. 2010. Secondary necrosis: the natural outcome of the complete apoptotic program. Febs Lett, 584, 4491–9.

Stirling, D. R., Swain-Bowden, M. J., Lucas, A. M., Carpenter, A. E., Cimini, B. A. & Goodman, A. 2021. CellProfiler 4: improvements in speed, utility and usability. Bmc Bioinformatics, 22, 433.

Stöhr, D., Schmid, J. O., Beigl, T. B., Mack, A., Maichl, D. S., Cao, K., Budai, B., Fullstone, G., Kontermann, R. E., Mürdter, T. E., Tait, S. W. G., Hagenlocher, C., Pollak, N., Scheurich, P. & Rehm, M. 2020. Stress-induced TRAILR2 expression overcomes TRAIL resistance in cancer cell spheroids. Cell Death Differ, 27, 3037–3052.

Tsuchiya, K., Nakajima, S., Hosojima, S., Thi Nguyen, D., Hattori, T., Manh Le, T., Hori, O., Mahib, M. R., Yamaguchi, Y., Miura, M., Kinoshita, T., Kushiyama, H., Sakurai, M., Shiroishi, T. & Suda, T. 2019. Caspase-1 initiates apoptosis in the absence of gasdermin D. Nat Commun, 10, 2091.

Webb, J. L. 1963. Enzyme and metabolic inhibitors, New York, Academic Press.

Yang, W. S., Sriramaratnam, R., Welsch, M. E., Shimada, K., Skouta, R., Viswanathan, V. S., Cheah, J. H., Clemons, P. A., Shamji, A. F., Clish, C. B., Brown, L. M., Girotti, A. W., Cornish, V. W., Schreiber, S. L. & Stockwell, B. R. 2014. Regulation of ferroptotic cancer cell death by GPX4. Cell, 156, 317–331.

Yang, W. S. & Stockwell, B. R. 2008. Synthetic lethal screening identifies compounds activating iron-dependent, nonapoptotic cell death in oncogenic-Ras-Harboring cancer cells. Chem Biol, 15, 234–45.

Yu, X., Wang, Y., Tan, J., Li, Y., Yang, P., Liu, X., Lai, J., Zhang, Y., Cai, L., Gu, Y., Xu, L. & Li, Y. 2024. Inhibition of NRF2 enhances the acute myeloid leukemia cell death induced by venetoclax via the ferroptosis pathway. Cell Death Discov, 10, 35.

Yuan, J. & Ofengeim, D. 2024. A guide to cell death pathways. Nat Rev Mol Cell Biol, 25, 379–395.

